# Integration of eye-centered and landmark-centered codes in frontal eye field gaze responses

**DOI:** 10.1101/791236

**Authors:** Vishal Bharmauria, Amirsaman Sajad, Jirui Li, Xiaogang Yan, Hongying Wang, J. Douglas Crawford

**Author notes:** Corresponding author: Dr. John Douglas Crawford, Departments of Psychology, Biology and Kinesiology & Health Sciences York University, Toronto, Canada, Centre for Vision Research, Room 0009A LAS, 4700 Keele Street, Toronto, Ontario, M3J 1P3.

## Abstract

The visual system is thought to separate egocentric and allocentric representations, but behavioral experiments show that these codes are optimally integrated to influence goal-directed movements. To test if frontal cortex participates in this integration process, we recorded primate frontal eye field (FEF) activity during a cue-conflict memory delay saccade task. To dissociate egocentric and allocentric coordinates, we surreptitiously shifted a visual landmark during the delay period, causing saccades to deviate by 37% in the same direction. To assess the cellular mechanisms, we fit neural response fields against an egocentric (eye centered target-to-gaze) continuum, and an allocentric shift (eye-to-landmark centered) continuum. Initial visual responses best fit target position. Motor responses (after the landmark shift) predicted future gaze position but embedded within the motor code was a 29% shift toward allocentric coordinates. This shift appeared transiently in memory-related visuomotor activity, and then reappeared in motor activity before saccades. Notably, fits along the egocentric and allocentric shift continua were initially independent, but became correlated just before the motor burst. Overall, these results implicate frontal cortex in the integration of egocentric and allocentric visual information for goal-directed action, and demonstrate the cell-specific, temporal progression of signal multiplexing for this process in the gaze system.

## INTRODUCTION

In our daily interactions with the surrounding world, the visual system relies on two ways of coding the relative location of visual objects: egocentric (e.g., object location relative to the eye) and allocentric (e.g., object location relative to an independent landmark) (Schenk 2006; Ball et al. 2009; Chen et al. 2011). For example, each word on this page stimulates the visual system in retinal coordinates, but the surrounding page borders provide higher level allocentric cues to remember their location. Neuropsychological and neuroimaging studies have suggested that these two types of visuospatial coding are segregated at an early stage of visual processing, generally associating egocentric transformations with ‘dorsal stream’ parietal cortex, and allocentric representations with ‘ventral stream’ temporal cortex (Goodale and Haffenden 1998; Milner and Goodale 2006; Byrne and Crawford 2010; Chen et al. 2014; Filimon 2015). The dual influence of such visual cues on cognition and behavior has been studied extensively in the context of perception (Murphy 1998; Kudoh 2005), memory (Chum et al. 2007; Coluccia et al. 2007; Byrne and Crawford 2010), and navigation (Colby 1998; Mou et al. 2006; Byrne et al. 2007; Wallach et al. 2018). However, it is not known how allocentric landmark cues are integrated into the egocentric motor commands for goal-directed action (Thaler and Goodale 2011; Chen et al. 2014; Fiehler et al. 2014; Chen, Monaco, et al. 2018). Understanding this is important for a general understanding of how visual cues are integrated and multiplexed for goal-directed action, and may ultimately help understand how the brain might compensate for loss of one of these visual coding mechanisms due to injury or disease (Murphy 1998; Schenk 2006).

Recent human psychophysics experiments — in which visual landmarks were shifted during the interval between seeing and acting — suggest that sensorimotor systems optimally integrate these cues (Byrne and Crawford 2010; Li et al. 2017). The weighting of allocentric cues is typically in the range of 30-50 % (Neggers et al. 2005; Byrne and Crawford 2010; Fiehler et al. 2014; Klinghammer et al. 2017a; Li et al. 2017), depending on factors such as perceived reliability of the cue, number of cues, predictability, aging and others (Bridgeman et al. 1997; Lemay et al. 2004; Neely et al. 2008; Byrne and Crawford 2010). Although there are several high-level theories of how such integration might occur (Byrne et al. 2007; Körding et al. 2007; Byrne and Crawford 2010; Mutluturk and Boduroglu 2014; Filimon 2015; Lew and Vul 2015; Klinghammer et al. 2017a; Aagten-Murphy and Bays 2019), the neural mechanisms were not tested. A recent neuroimaging study found that certain areas of frontal and parietal cortex are active *when* participants use allocentric information to guide action (Chen, Monaco, et al. 2018), but neurophysiological recordings would be necessary to directly show *if* and *how* integration is occurring in these sites. In contrast to the many neurophysiological studies of allocentric coding for memory and navigation (O’Keefe 1976; Rolls 1999), very few studies have investigated the neurophysiology of allocentric coding in visual and visuomotor systems. For example, the influence of allocentric landmarks has been studied in neurons of cingulate cortex (Dean and Platt 2006), entorhinal cortex (Meister and Buffalo 2018), and precuneus (Uchimura, M; Kumano, H; Kitazawa 2017). Further, superior colliculus discharge is influenced by surrounding landmarks (Edelman and Goldberg 2003), and neurons in the supplementary eye fields (SEF) have been shown to encode object-centered (one part of an object relative to another) saccades (Olson and Gettner 1995; Tremblay et al. 2002). To our knowledge, the currently available literature does not describe how egocentric and landmark-centered signals are combined at the neuronal level for goal-directed action.

Based on this body of literature, we hypothesized that prefrontal cortex might play a special role in the re-integration of allocentric information into the egocentric visuomotor commands. Specifically, we hypothesized that the frontal eye field (FEF) should play a prominent role for this function in gaze control. The FEF is a saccade-related premotor cortical area with well-organized visual and motor response fields (RF) (Bruce and Goldberg 1985; Schall 2015) related to target selection (Schall et al. 1995; Lee and Keller 2008; Sajad et al. 2015; Raghavan and Joshua 2017), saccades (Robinson and Fuchs 1969; Guitton and Mandl 1978; Bruce and Goldberg 1985), has inputs from areas such as the SEF and lateral intraparietal cortex (LIP) (Munoz 2002), and also has direct outputs to the brainstem oculomotor and head movement control centers (Sommer and Wurtz 2000; Isa and Sasaki 2002). Thus, one would expect any cortical signal that integrates egocentric and allocentric information to be present at this point in the system. We recently found that (in the absence of landmarks) the FEF produces a target-to-gaze coding transformation in egocentric coordinates (Sajad et al. 2015, 2016). Based on this, one might hypothesize that an allocentric influence should occur within and between these codes.

To test this hypothesis, we trained two rhesus macaques to perform a memory delay, cue-conflict gaze task similar to that used in previous behavioral studies of egocentric-allocentric integration (Fig. 1A) (Li et al. 2017). We then used this task to explore visual, memory, and motor RFs in the FEF, employing a model-fitting procedure (Keith et al. 2009; DeSouza et al. 2011; Sadeh et al. 2015; Sajad et al. 2015) to characterize both the egocentric and allocentric codes used in neurons, i.e., if their activity only predicts future gaze position in an egocentric frame or if it also predicts the influence of a landmark shift on gaze behavior. We did these experiments in head-unrestrained conditions in order to first confirm the optimal egocentric coordinate frame (Sajad et al. 2015), and then extended this approach to fitting landmark-centered models to test the following questions: 1) if a landmark shift influences spatial coding between visual and motor responses, 2) when this influence occurs (i.e., immediately, gradually, or delayed), and finally 3) how this allocentric influence is integrated within the egocentric code (i.e., independently, or integrated through some specific multiplexing mechanism). We found that the FEF does play an important role in this process, progressively integrating allocentric visual information into the egocentric gaze code through different cell types to yield an integrated cortical motor gaze command.

**Fig. 1.**
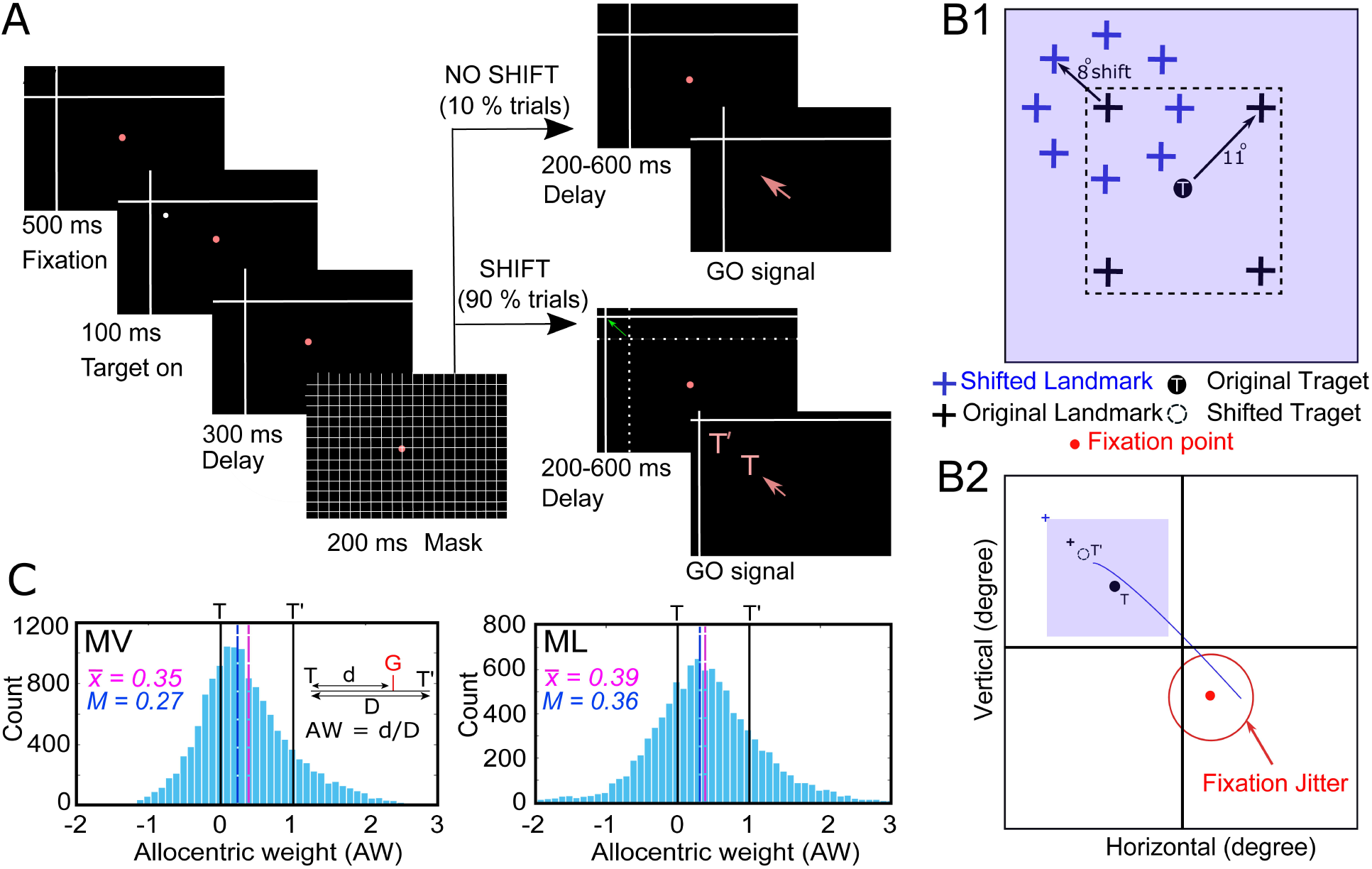
Experimental paradigm and behavioral results. **(A)** Cue-conflict task and its time course. The monkey fixated on a red dot for 500 ms while a landmark (L, intersecting lines) was already present on the screen. Next, a target (white dot) was flashed for 100 ms, and after a delay of 300 ms, a grid-like mask was presented (200 ms). After a second memory delay (200-600 ms), the monkey was cued (disappearance of the red fixation dot, i.e., go signal) to saccade toward the remembered location of target (T’, virtually shifted remembered target, i.e. the virtual target location if entirely relying on the shifted landmark) either in presence of a shifted landmark (L’, indicated by green arrow) or in absence of it. The red arrow corresponds to the head-unrestrained gaze shift toward the memorized location of the target (T = original location of target; T’ = virtually shifted target location). To avoid biasing the behavior, animals were rewarded for placing gaze anywhere within 8-12° radius of the reward window centered on T. **(B1)** The visual landmark was presented in one of the four oblique directions 11° from the target, and it shifted 8° from its initial position in one of the eight radial directions. An example of all (four) possible landmark (black cross) locations for an example target (black dot, T). After the mask offset, the landmark shifted (by 8°, blue cross represents the shifted landmark) in one of eight radial directions around the initial presentation of the landmark. **(B2)** An example of a gaze shift (blue curve) from the initial fixation point (red dot) toward the virtually shifted target (broken circle, T’), in relation to the shifted landmark (black cross, L’). **(C)** Distribution (X-axis) of allocentric weight (AW) for monkey V (MV, left) and monkey L (ML, right) plotted against the number of trials (Y-axis) collapsed across all the analyzed trials for all spatially tuned neurons. The mean AW (vertical pink line) for MV and ML was 0.35 and 0.39 (mean = 0.37) respectively. **Note:** the ‘no shift’ condition only included 10 % of the trials whereas the ‘shift’ condition included 90 % of trials. However, the AW was computed on the shifted condition only. The ‘no-shift’ condition was included to test all the range of the shifts.

## MATERIALS AND METHODS

### Surgical Procedures and recordings of 3D Gaze, Eye, and Head

All experimental protocols were approved by the York University Animal Care Committee and were in accordance with the guidelines of Canadian Council on Animal Care on the use of laboratory animals. Neural data were recorded from two female *Macaca mulatta* monkeys (MV and ML). Animals were prepared for 2D and 3D eye-movement and electrophysiological recordings. Surgeries were performed as done previously (Crawford et al. 1999; Klier et al. 2003). The recording chamber was implanted and was centered in stereotaxic coordinates at 25 mm anterior and 19 mm lateral for both monkeys. A craniotomy of 19 mm (diameter) covering the base of the chamber (attached over the trephination with dental acrylic) allowed access to the right FEF. Two 5-mm-diameter (a 2D and a 3D) sclera search coils were implanted in the left eye of both animals. During experiments, animals were seated within a custom-made primate chair adapted to allow free movements of the head at the center of three mutually orthogonal magnetic fields (Crawford et al. 1999). This allowed us to record 3D eye movements (i.e., gaze) and orientation (horizontal, vertical, and torsional components of orientation of the eye relative to space). Two orthogonal coils that allowed similar recordings of the head orientation in space were also mounted during the experiment. These values allowed us to compute other variables such as the eye orientation relative to the head, eye, and head velocities, and accelerations (Crawford et al. 1999).

### Basic Behavioral Paradigm

Using laser projections, the visual stimuli were presented on a flat screen, placed 80 cm away from the animal (Fig. 1A). To analyze visual (to target presentation) and movement-related (gaze shift onset) responses in the FEF separately, monkeys were trained to perform a standard memory-guided gaze task to a remembered target in relation to a visual landmark (two intersecting lines, acting as an allocentric landmark), thus imposing a temporal delay between presentation of the target and movement initiation of the eye. A trial began by the animal fixating on a red dot, placed centrally, for 500 ms in the presence of the landmark. Then a visual target (T, white dot) was briefly flashed for 100 ms, followed by a delay (300 ms), a grid-like mask (200 ms, this occludes past visual traces, current and future landmark) and a memory delay (200-600 ms). As the initial fixation target extinguished, the monkey was cued to saccade head-unrestrained toward the remembered location of the target either in the presence of a shifted landmark (90 % of trials, shifted by 8° in one of the eight radial directions) or in absence of it (10 %, no shift condition, i.e. landmark appeared in the same position as before mask). The task is detailed in Fig 1B. In the shift condition, the target virtually shifted (T’) in relation to the landmark, and the monkey was rewarded a drop of water to place his gaze (G) within 8-12° radius of the original target (so that the animals were rewarded if they looked at T, at T’, or anywhere in between). This large reward window allowed for the variability of memory-guided gaze shifts and accumulation of errors (Gnadt et al. 1991; White et al. 1994; Sajad et al. 2015), which formed the logical premise of our analysis (see below). The influence of landmark shift on behavior (Fig. 1C) was computed as allocentric weight (AW) as follows:

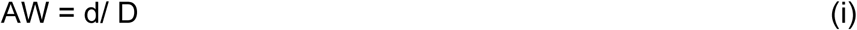

where *AW* is allocentric weight; *d* is the projection of TG onto TT’, and *D* is the magnitude of TT’. The output values (AW) are between 0 and 1 mainly, where 0 corresponds to absolute egocentric coding and 1 refers to absolute allocentric coding, where the gaze landed on T’, completely following the shift in the cue location (Byrne and Crawford 2010; Li et al. 2017).

### Electrophysiological Recordings and sampling of gaze, eye and head

The extracellular activity from FEF neurons was recorded using tungsten microelectrodes (0.2–2.0 mΩ impedance, FHC Inc.) and it was digitized, amplified, filtered, and stored for offline spike sorting using template matching and further applying principal component analysis on the separated clusters with Plexon MAP System. The recorded sites of FEF (in head-restrained conditions) were further confirmed by employing a low-threshold (50 μA) electrical microstimulation as defined previously (Bruce et al. 1985). An overlapped pictorial of the sites from both animals is presented in Fig. 2A-B (ML in Blue and MV in red). We recorded an average of 331 ± 156 (mean ± SD) trials/neuron. Only the sites within the arcuate sulcus were carried forward for analysis. Mostly, neurons were searched for while the animal was freely (head-unrestrained) scanning the environment. Once a neuron had reliable spiking activity, the experiment started. A neuron’s RF (visual or motor) was characterized while the monkey performed the task and made memory-guided gaze shifts from a fixation point to randomly presented targets (presented one by one, generally contralateral to the recording site) within an array (varying between 4 × 4 to 7 × 7 depending on size and shape of the RF) of targets (5 –10° apart). Initial fixation positions were also jittered within a 7-12° window that accounted for the increase in the variability of initial 3D gaze, eye, head distributions and displacements for our analysis, thus allowing effector separation. The position of eye and head orientations (in space) were recorded with a sampling rate of 1000 Hz. For movement analysis, the gaze shift onset (eye movement in space) was marked at the time when the gaze velocity was more than 50°/s and the gaze offset was selected as a time point when the velocity declined below 30°/s. The movement of the head was marked from the gaze shit onset till the time point at which velocity of the head declined below 15°/s.

**Fig. 2.**
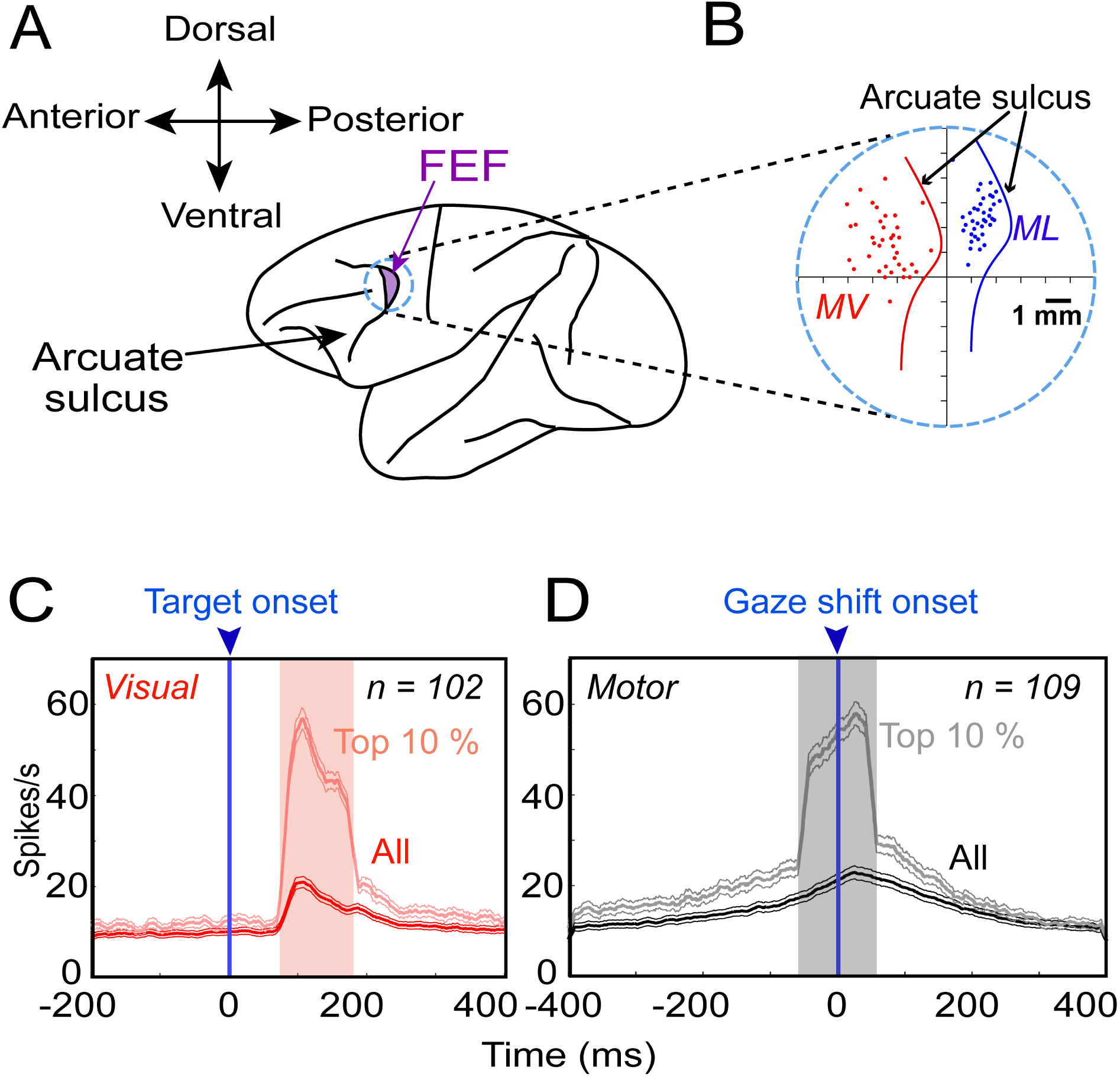
FEF recordings. **(A)** The purple patch shows the location of the FEF inside the arcuate sulcus and the circle (although does not correspond to original dimensions of chamber) represents the chamber. **(B)** An overlapped zoomed-in section of the chamber and the recording sites (dots) confirmed with 50 µA electrical stimulation from MV (red) and ML (blue). **(C)** Average (± 95 % confidence interval) of the spike-density plots (dark red: all trials; light red: top 10% best trials most likely corresponding to the hot spot of response fields, RFs) of every neuron in visual population, aligned to the target onset (blue arrow). **(D)** same as **C,** but for motor neurons in relation to the gaze shift onset (blue arrow). **Note:** the shaded area corresponds to the 100-ms sampling window.

### Data Inclusion Criteria, sampling window and neuronal classification

Only clearly isolated and task-modulated neurons were analyzed, i.e., cells with clear visual activity and/or with perisaccadic movement activity (Fig. 2C-D). Neurons with post-saccadic (activity after gaze shift onset), or delay activity were excluded. Only trials where monkeys landed their gaze within the acceptance window were included. For neural activity analysis, the sampling window for “visual epoch” was characterized as a fixed 100-ms window of 80 – 180 ms after the target onset and the “movement period” was defined as a perisaccadic 100-ms (−50 – +50 ms relative to gaze shift onset, because it corresponded to high-frequency perisaccadic burst) period (Sajad et al. 2015). This provided a good signal-to-noise ratio for neuronal analysis, and most likely represented the time-period in which gaze shifts were influenced by FEF activity. Further, neurons were characterized based on trials that showed the top 10% of spiking activity in the fixed sampling window, called “Vis10%” for the visual activity and “Mov10%” for the movement activity. These top 10 % trials approximately corresponded to the peak response within the RF. To quantify the relative strength of visual versus the motor activity, a visuo-movement index (VMI) for each cell was computed as the difference of Vis10% and Mov10% divided by their sum, after subtracting off their trial-matched baseline activity.

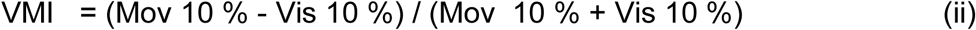

The VMI values fall between – 1 and + 1, where −1 represents a purely visual neuron and + 1 corresponds to a purely motor neuron.

### Spatial Models considered in Preliminary Analysis

In an exhaustive preliminary analysis, we tested if 1) the presence of a visual landmark fundamentally changed the FEF codes reported in our previous studies (Sajad et al. 2015, 2016) and 2) if we could narrow the focus of our analysis in the current study. First, we fitted (using Matlab 2015a) visual and motor RFs against the eleven egocentric canonical models **(Supplementary Fig. S1A)** that were previously analyzed (Sajad et al. 2015) based on the different spatial parameters (most importantly target, T, and final gaze position, G) involved in a memory-guided task. These models are as follows: dH, final minus initial head orientation in space coordinates; dE: final minus initial eye orientation in head coordinates; dG, final minus initial gaze position in space coordinates; Hs, final head position in space coordinates; Eh, final eye position in head coordinates; Gs, Gh, Ge — Gaze in space, head and eye coordinates respectively; Ts, Th, Te — Target in space, head and eye coordinates respectively. A further detail of these models is provided in the previous analysis (Sajad et al. 2015). Since this task involved an allocentric landmark, we then added another eight models of target coding based on the initial and shifted landmark location. **Supplementary Fig. S1B** shows different models considered in this study (in addition to the preliminary analysis on 11 egocentric models) and how we derived them. In the allocentric analysis, we retained three egocentric models (Ts, Te and Ge) for comparison. The allocentric models are as follows: Ls, Landmark within the space coordinates; L’s, Shifted landmark within the space coordinates; Le, Landmark within eye coordinates; L’e, Shifted landmark within eye coordinates; T’s, Shifted target in space coordinates; T’e, Shifted target in eye coordinates; TL, Target relative to landmark; TL’, target relative to shifted landmark. Note: the prime (‘) here means positions that are related to the new (shifted) location of the landmark. As summarized below in the Results section and in supplementary figures **(S2-S5),** this preliminary analysis led to the conclusion that the Te and Ge models continued to dominate visual and motor activity respectively in this task (Sajad et al. 2015) which led us to focus our analysis on testing the following, more specific spatial models.

### Intermediate spatial models used in main analysis

In our previous studies, we found that FEF responses did not necessarily fit best against exact spatial models like Te or Ge, but may instead fit best against models intermediate between those canonical types (Fig. 3A, lower left) (DeSouza et al. 2011; Sadeh et al. 2015; Sajad et al. 2015, 2016). Different intermediate model continua were analyzed as in the previous study (Sajad et al. 2015) involving egocentric models and it was revealed that the Te-Ge (Target in eye coordinates to Gaze in eye coordinates) was the most relevant and most of the transformations occurred along this continuum (see results). Based on this previous finding and our preliminary model-fitting analysis (summarized below in the Results section and Supplementary Figures), we retained this continuum (Te to Ge) to quantify the FEF egocentric transformation (Fig. 3A, lower left).

**Fig. 3.**
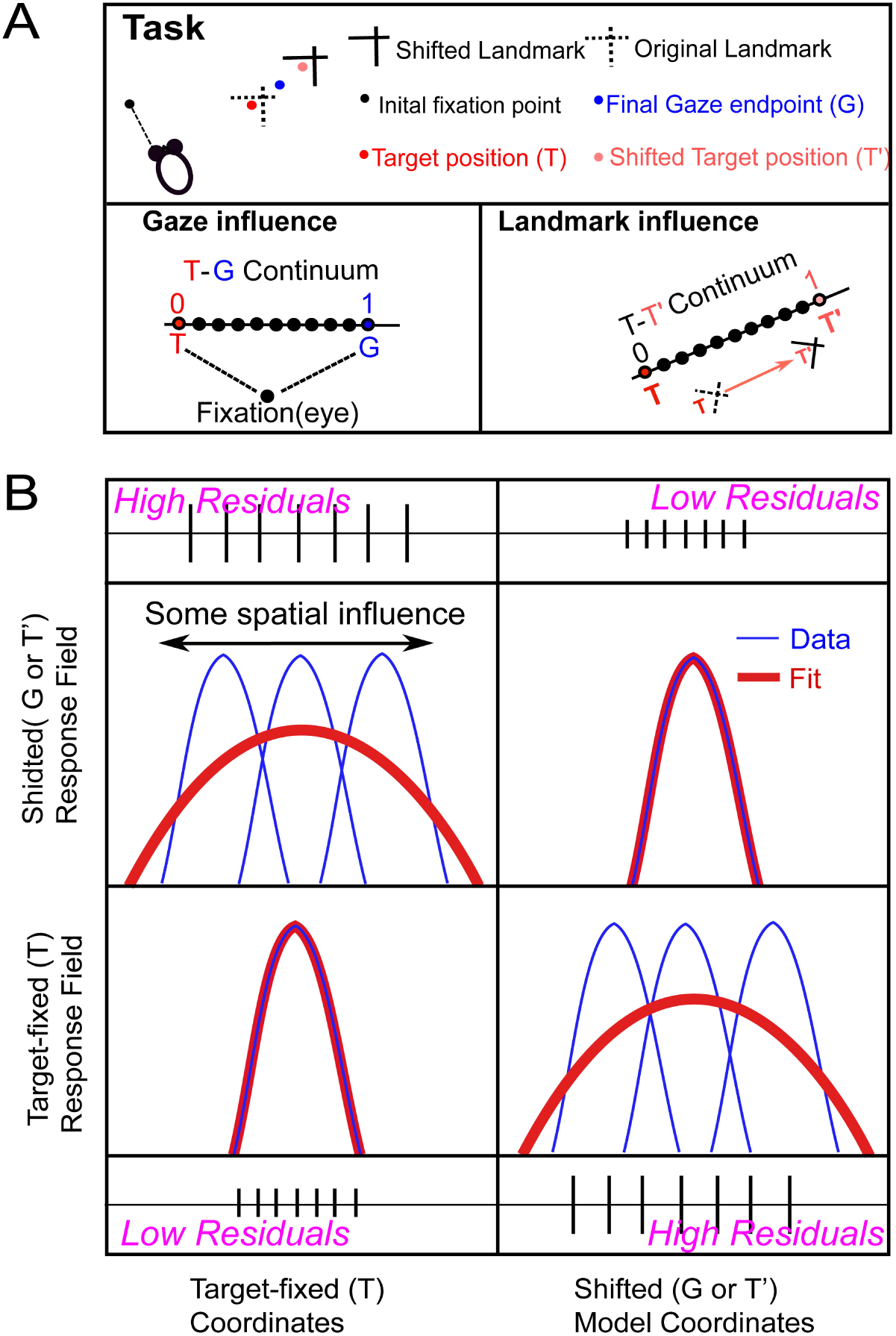
Schematic representation of spatial parameters and model fitting procedure. **(A)** Schematic illustration of spatial parameter for a single trial. The projections of initial fixation/gaze (black), and final gaze (blue). The red dot represents the target (T) location relative to the original landmark (L) position (broken intersecting lines), whereas the lighter dot (red) corresponds to the virtually shifted target (T’) relative to the shifted landmark (L’) position (solid intersecting lines). Notice that the final gaze position lands between the T and T’ (i.e., shifted toward the shifted landmark). In head-unrestrained conditions, one can encode the positions in egocentric coordinates (eye, head or body) (Sajad et al. 2015). Here, we plotted two continua: a T-G spatial continuum (egocentric, i.e., we treated these codes as a continuous spatiotemporal variable by dividing the space between T and G into ten equal steps) to compute the gaze influence and a T-T’ spatial continuum (egocentric to allocentric) based on the landmark shift to compute the landmark influence on the neuronal activity. **(B)** A schematic of the logic of RF analysis. X-axis depicts the target-fixed coordinates, and the Y-axis represents the corresponding activity to the target. If the activity to a fixed target location is fixed (bottom left), then the data (blue) would fit (red) better on that, thus yielding lower residuals in comparison with when the activity for the target is shifted or broadly tuned, thus leading to higher residuals (top left). Similarly, if the activity is fixed in shifted coordinates, this would lead to lower residuals and vice versa.

Further, based on our hypotheses, the shifted landmark would influence the gaze to remembered location. In order to isolate the influence of the landmark shift, we constructed (Fig. 3A, right) another continuum (T-T’) between Te and T’e (shifted target in eye coordinates) with ten steps (intermediary frames) in between, and additional steps on either sides (10 beyond Te and 10 beyond T’e), i.e., the spatial continuum from eye-centered to landmark centered target coding (Fig. 3A, right), similar to the allocentric weight score used to describe the behavior (Fig. 1C). The T-T’ comparison will test whether FEF code is purely based on the ego-centric information (T), allocentric information (T’), or contains the integrated allocentric + egocentric information that eventually drives the behavior.

Henceforth, we will refer to these as the egocentric (T-G) and allocentric shift (T-T’) continua. These additional ten steps on either side were accommodated 1) to allow for the possibility that neurons can encode for abstract spatial codes outside the canonical models and 2) to avoid misleading edge effects, otherwise the preferred spatial models would cluster around the canonical models. Thus, these two continua (T-G and T-T’) proved to be the most useful in describing the FEF transformations that occurred in the current task.

### Fitting Neural Response Fields Against Spatial Models

To test different models as described above, they must be spatially separable (Keith et al. 2009; Sajad et al. 2015). This spatial separation was provided by natural (untrained) variations in the monkeys’ behavior. For example, the variability in memory-guided gaze shifts allowed us to distinguish target coding form the gaze coding, the initial eye and head movement allowed us to separate different egocentric reference frames and variable movements of the eye and head for a gaze shift allowed us to distinguish different effectors. In contrast to decoding methods that typically test if a spatial parameter is implicitly coded somehow in patterns of population activity (Bremmer et al. 2016; Brandman et al. 2017), our method directly tests which spatial model best predicts activity in spatially tuned neurons and populations. The schematic in Fig. 3B describes the logic of RF fitting in different reference frames. Essentially, RF data plotted in the correct spatial frame will yield the lowest residuals (errors between the fit and data points) compared with other models. Suppose that a neural response encodes target position (T) in eye-coordinates (Fig. 3B, lower row quadrants) — in this case, if a fit is computed to its RF in the same coordinate system (T), then the fit will match the data and yield low residuals (Fig. 3B, lower-left quadrant). However, if the RF fits were computed in some other coordinate system, like future gaze position (G, which often shifts from T due to variable motor errors (Sajad et al. 2015)) or landmark-centered target position (T’, which shifts in our cue-conflict task), then the fit will not match the RF and yield higher residuals (Fig. 3B, lower-right quadrant). Conversely, if the actual neural RF is organized relative to G or T’, then fits in these coordinates should give lower residuals than fits in the unshifted T coordinates (Fig. 3B, upper row quadrants). Likewise, if the actual RF was organized at some intermediate point between T and G or T and T’ (Fig. 3 B), this point will yield least residuals (Sajad et al. 2015).

In practice, we used a non-parametric fitting technique to characterize neural activity relative to spatial location and varied the spatial bandwidth of the fit to match any RF size, shape, or contour (Keith et al. 2009). To test between various spatial models, we used Predicted Residual Error Some of Squares (PRESS) statistics. To independently calculate the residual for a trial, its actual activity was subtracted from the corresponding point on a fit made to all the other trials (similar to cross-validation). Note that if the physical shift in spatial positions between two models only results in a systematic shift (direction and amount), this would appear as an overall shift or expansion in RF and our statistical modeling method would not be able to dissociate between the two models as they would yield virtually indistinguishable residuals. But since in our study, the distribution of relative positions between different spatial models also has a non-systematic variable component (e.g., variable errors in gaze endpoint, or unpredictable landmark shifts) the RFs generally stayed in the same location, but the spatial models were dissociable based on the residual analysis.

The visual and movement RFs of neurons were plotted in the coordinates of all the tested canonical (and intermediate) models. For mapping of visual response in egocentric coordinates, we used eye and head orientations at the time of the target presentation, whereas for motor RF mappings we used behavioral measurements at the time that the gaze shift began (Keith et al. 2009; DeSouza et al. 2011; Sajad et al. 2015). Similarly, for the allocentric models, we used the initial location of the landmark and the shifted location of the landmark to plot our data. As we did not know the shape and size of RF a priori and the spatial distribution of the data was different for every model (e.g. the models would have a smaller range for head than the eye models), we plotted the non-parametric fits with kernel bandwidths ranging from 2-25°, thus ensuring that we were not biasing our fits toward a particular size and spatial distribution. Across all tested models with different non-parametric fits, we calculated the PRESS residuals to infer the best model for that neural activity (that yielded the least PRESS residuals). The mean PRESS residuals of the best model were then statistically compared with that of the other models at the same kernel bandwidth using a two-tailed Brown-Forsythe test (Keith et al. 2009a). Finally, we did a statistical analysis (Brown-Forsythe) to compare the mean PRESS residuals of different spatial models for population analysis (Keith et al. 2009a; DeSouza et al. 2011). A similar procedure was employed to compute the best fits for the intermediate model continua.

### Testing for Spatial Tuning

The method described above assumes that neural activity is organized into spatially tuned RFs. This does not mean that other neurons do not contribute to the overall population code (Goris et al. 2014; Bharmauria et al. 2016; Leavitt et al. 2017; Chaplin et al. 2018; Zylberberg 2018; Pruszynski and Zylberberg 2019) but rather that only these neurons can be explicitly tested. Here, we tested for the spatial tuning at the single cell. We compared the spatial selectivity of the neuron by randomly shuffling the firing rate data over the position data obtained from the best-fit model (100 times to obtain random 100 RFs). The distribution of the mean PRESS residuals of the 100 randomly generated RFs were then statistically compared to the mean PRESS residual (PRESS_best-fit_) of the best-fit model (unshuffled, original data). If the mean PRESS of the best-fit model fell outside of the 95% confidence interval of the shuffled mean PRESS distribution, then the neuron’s activity was categorized as spatially selective. At the level of the population analysis, some neurons exhibited spatial tuning at certain time-steps whereas others did not because of low signal to noise ratio. Therefore, we eliminated time steps where the mean spatial coherence (goodness of fit) of the population was statistically indistinguishable from the baseline (before target) as there was no task-relevant information at this stage and thus had no spatial tuning. For this, we defined an index for spatial tuning, Coherence Index (CI) for a single neuron, which was computed as (Sajad et al. 2016):

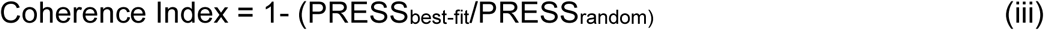

If the PRESS_best-fit_ was similar to PRESS_random_ then the CI would be approximately 0, and if the best-fit model is a perfect fit (i.e., PRESS_best-fit_ = 0), the CI would be 1. Only neuronal activity that showed significant spatial tuning, as indicated by these methods, was included in the analysis presented below.

### Time-normalization for spatiotemporal analysis

The temporal structure of the task was such that the time between target onset, and the time of gaze onset were not equal across trials. To correctly account for transformations between these events, it is essential to account for this variability. We tackled this by normalizing the duration of each trial. Accordingly, the activity across different trials (between the events of interest — from 80 ms after the visual stimulus onset until gaze shift onset for whole population in Figure 7, and from mask off until gaze shift onset for subpopulations in Figure 8) was divided into an equal number of half overlapping bins. For the whole population, the activity from different trials was divided into 14 half overlapping equal bins (ranging from 112.26 – 178.93 ms depending on the trial) and into 7 half overlapping bins for the individual populations. The rationale behind the choice for bin number was to ensure the sampling time window is wide enough, and thus, robust for differences due to the stochastic nature of spiking activity. During the whole population analysis, the normalization procedure also caused the duration of the mask to be convolved in time, on average from step 4.69 to step 7.63. For subpopulation analysis, the bin size ranged from 80.5 – 180.5 ms. In this paradigm with variable delay activity, aligning trials in a standard fashion (relative to visual stimulus onset or saccade onset) would result in the loss and/or mixing of neural activities across trials. Thus, this would not allow us to track spatial coding through the entire trial across all trials. Therefore, the time-normalization was done because of the different delay period in our case and the details of time-normalization are provided in the previous study (Sajad et al. 2016). Again, only spatially tuned data were included at every time-step to compute the spatiotemporal progress along the T-G and T-T’ continua. Specifically, for the spatiotemporal analysis, the neural firing rate (in spikes/second; the number of spikes divided by the sampling interval for each trial) was sampled at 14 half overlapping windows from these time-normalized data. These sampling window numbers were chosen based on the approximate ratio of the duration of the visual response to delay period to movement response (Sajad et al. 2016). The final (14th) time step in Figure 7 corresponded to some part of the perisaccadic sampling window as mentioned above. Therefore, this time-normalization of neural activity across trials allowed us to consider the entire sequence of visual–memory–motor responses as a continuum.

## RESULTS

### Influence of the landmark shift on behavior

As noted above, we used a cue-conflict task, where a prominent visual landmark shifted surreptitiously during the memory delay between the target presentation and the cue for a gaze saccade (Fig. 1A). All possible landmark (black cross) locations in relation to an example target (black dot, T) are shown in Fig. 1B1. The landmark was located 11° from the target in oblique directions. After the mask offset, the landmark shifted (by 8°, blue cross represents the shifted landmark) in one of eight radial directions around the initial presentation of the landmark. Fig. 1B2 shows an example of a gaze shift (blue curve) from the initial fixation point (red dot) toward T’ (broken circle), in relation to L’ (blue cross). The influence of the landmark shift on gaze behavior was described previously (Li et al. 2017), and we confirmed this in the current dataset (i.e., in the behavioral data recorded in conjunction with the neural data analyzed below). To do this, we computed the component of gaze end points along the axis between the target and the direction of cue shift, calculated as allocentric weight (AW); If AW = 0, that means the shifted landmark has no influence on that gaze and if AW = 1, a complete influence of landmark on gaze; see methods). This resulted in a large scatter of gaze end points along the axis of landmark shift (Fig. 1C), but there was a highly significant (p < 0.01, one sampled Wilcoxon signed rank test) tendency to shift away from the original target in the direction of the cue shift, with a mean allocentric weight of 0.35 in monkey V and 0.39 in monkey L. This agrees well with previous behavioral studies (Neggers et al. 2005; Byrne and Crawford 2010; Fiehler et al. 2014), including ours from the same animals (Li et al. 2017).

### Neural Recordings: General Observations and Neuronal Classification

To examine if and how the behavior described above was encoded in FEF activity, we simultaneously probed the visual and motor RFs from over 250 FEF sites using tungsten microelectrodes while monkeys performed the above task (Fig. 2A-B). After applying our exclusion criteria to the neural data (see methods), we were left with 102 neurons with significantly spatially tuned visual responses and 109 neurons with significantly spatially tuned motor responses, which often overlapped in visuomotor neurons. The averaged spike density plots of all visual (red) and all motor (black) neurons, with the 100-ms sampling window (shaded areas) used for analysis are displayed in Fig. 2C and 2D respectively. Our non-parametric analysis along both continua (as described in methods in Fig. 3A-B) utilized all trials throughout each neuron’s RF (dark black and red lines), but we also show the top 10% responses during the analysis window (light red and gray for visual and motor responses respectively), generally corresponding to the RF ‘hot spot’ activity recorded and analyzed in many studies. Following standard practice in the field (Bruce et al. 1985; Sajad et al. 2015; Schall 2015; Khanna et al. 2019), we further divided these populations into visual (only visual responses, Fig. 4A), visuomotor (both visual and motor, Fig. 4B) and motor (only motor responses, Fig. 4C) neurons, using the same convention for top 10 % and all responses as above.

**Fig. 4.**
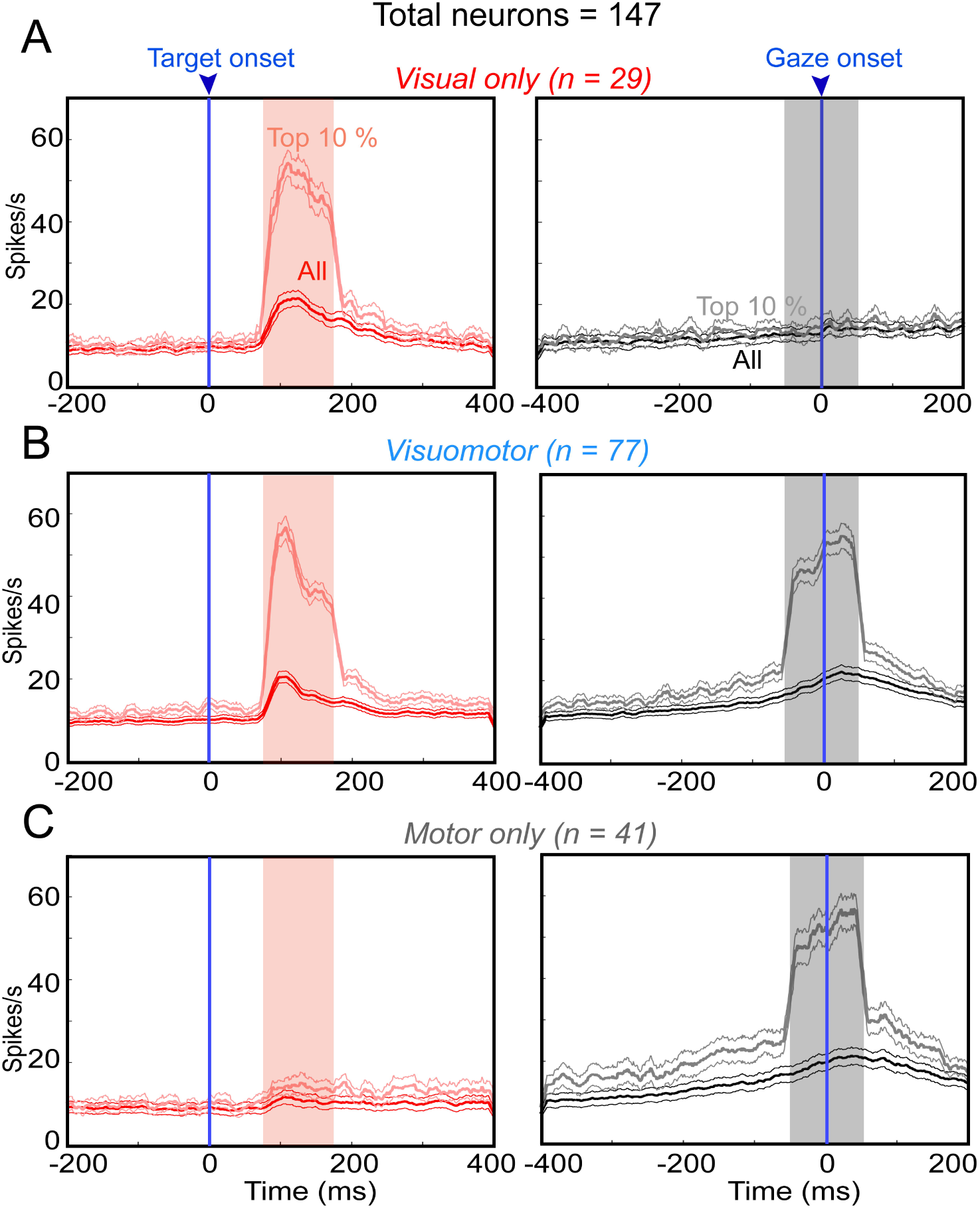
Spike density plot of different categories of neurons. **(A)** Averaged spike density (computed similarly as Figure 2) plots of visual only (V) neurons aligned to target on (blue line, left panel) and gaze shift onset (blue line, right panel). **Note:** V neurons respond (red shaded area, the red curve represents all responses and the light red curves corresponds to top 10 % responses) only to the target onset, whereas there is no response to gaze shift onset. **(B)** same as ***A*** but for visuomotor (VM) neurons. Note: the VM neurons respond to both target on and gaze shift onset. **(C)** same as ***A*** and ***B*** but for motor only (M) neurons. **Note:** M neurons have a pronounced response only to gaze shift onset. Red and grey shaded areas denote the sampling windows for RF analysis.

### Does the landmark fundamentally alter the FEF egocentric transformation?

As noted in the methods, we began our analysis of these neurons by testing if the visual landmark altered the codes that we previously observed in FEF visual (primarily target-in-eye coordinates) and motor (primarily gaze-in-eye coordinates) in the absence of a landmark (Sajad et al. 2015). Although this analysis was exhaustive, we only summarize it here because the results were very similar to the previous results. However, these results were important to establish the subsequent analyses.

We began this analysis by considering whether the addition of the landmark (and/or its shift) obviate the basic egocentric target-to-gaze transformation reported in our previous FEF studies (Sajad et al. 2015, 2016). Overall, we found that the T and G (in eye coordinates) models continued to dominate visual (**Supplementary Fig. S2 and S3)** and motor responses (**Supplementary Fig. S4 and S5)** respectively, i.e., they yielded lower residuals compared with residuals over all other egocentric models tested. This remained the case when we compared T and G to various possible ‘pure’ allocentric models, such as target relative to landmark in various coordinates, i.e., T and G were still dominant **(Supplementary Fig. S6).** Thus, in the presence of the landmark, FEF continued to be dominated by an eye-centered transformation (Fig. 5).

**Fig. 5.**
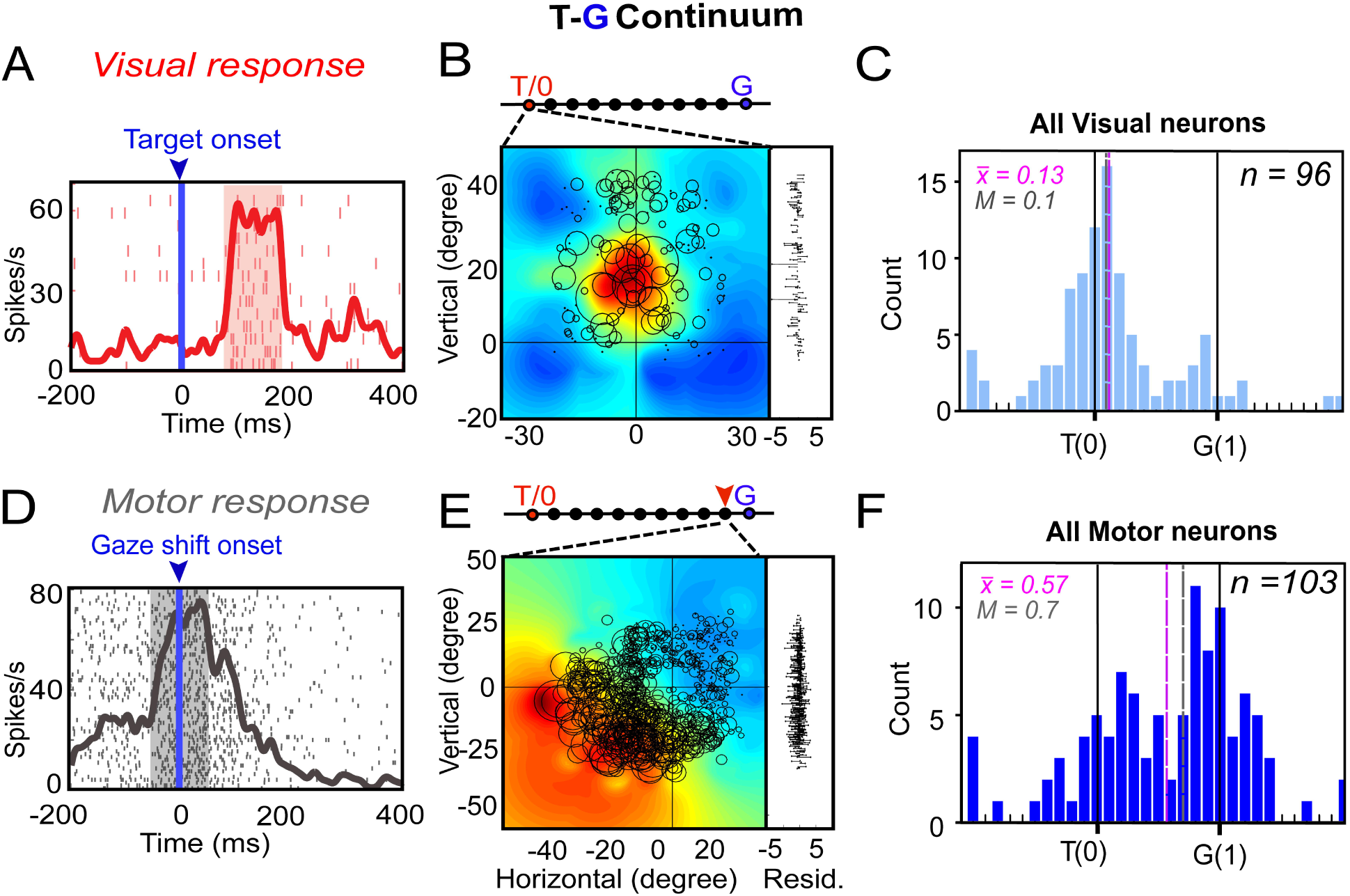
Egocentric Analysis (T-G continuum) of Visual and Motor Responses. **(A)** Raster/spike density plot of the neuron aligned to the target onset (blue arrow); the shaded red region represents the sampling window (80-180 ms) for response field (RF) analysis. **(B)** Representation of the RF using the eye-centered coordinates derived from the best-fit of the data along the T-G continuum (indicated by the red arrow on the scale above the RF). The diameter of the circles indicates the number of action potentials in the sampling window of each trial, whereas the color map represents a non-parametric fit to these data. For example, the red-hot region corresponds to the maximal activity and ‘hot spot’ of the RF. To the right of the plot are the residuals between the individual trial data and the RF fit. To independently compute the residual for a trial, its activity was subtracted from the corresponding point on the fit made to the rest of the trials. **(C)** Frequency distribution of all spatially tuned visual neurons (n = 96) along the T-G continuum with the best fit around T (mean = 0.13; median = 0.1). **(D)** Raster/spike density plot of a motor neuron aligned to the gaze shift onset (blue arrow); the shaded gray region represents the sampling window (–50-50 ms) for RF analysis. **(E)** An example of RF analysis of a motor response along the T-G continuum. RF representation along the T-G continuum, the RF fits best at the ninth step (red arrowhead) from T (one step closer to G). **(F)** Frequency distribution of all spatially tuned motor neurons (n = 103) along the T-G continuum with a significant influence of gaze (mean = 0.57; median = 0.7; p < 0.0001, one sampled Wilcoxon signed rank test). **Note:** the color bar indicates the magnitude of activity of neurons and corresponds to the size of circle in the RF map.

Fig. 5A-B shows an example analyses of a typical visually responsive neuron. Fig. 5A displays the raster/spike density plot of the visual neuron’s activity, aligned to the target onset (blue arrow, shaded area represents the 80-180 ms sampling window). Note that these data were collected in the presence of the landmark, but before the landmark shift. Fig. 5B shows the corresponding RF: the circles indicate the size and RF location of individual trial responses, and the background heat map shows the non-parametric fit to these data (where the red-hot blob indicates a typical ‘closed’ RF), and the corresponding residuals are plotted to the right. The location of best fit along the T-G continuum is indicated above the RFs. The best fit of the visual response was located exactly at T. This observation is documented for all neurons with significantly tuned visual responses (n = 96) in Fig. 5C. The distribution of best fits along the T-G continuum (Fig. 5C) peaked near T (mean = 0.13; median = 0.1). There was a slight but significant shift toward G due to a secondary peak near G — perhaps signifying that some of these responses already predicted future gaze position — but most units did not show this shift.

Fig. 5D shows the raster/spike density plot for a typical motor response, aligned to gaze shift onset. Fig. 5E shows the best-fit RF, in this case showing a typical ‘open’ RF, with activity that continues to grow with saccade size down-left. Note that these data are collected in the presence of the shifted landmark. In this case, along the T-G continuum, the RF was best located at the 9^th^ step from T (nearly at G, as indicated by the red arrow). This observation is documented across all neurons (n = 103) with significantly tuned motor responses in Fig. 5F. In contrast to the visual data, the motor response (Fig. 5F) exhibited a significant shift to the right toward G (mean = 0.57, median = 0.7; p < 0.0001, one sampled Wilcoxon signed rank test), with the major peak closer to G. In other words, most motor responses shifted along this eye-centered continuum to predict future gaze position. Comparing Fig. 5C and 5F, there was a significant shift between the visual and motor distributions along the T-G continuum (p < 0.0001; Mann-Whitney U-test). Further, as we reported previously (Sajad et al. 2015) this also occurred *within* cells with both visual and motor responses (**Supplementary Fig. S7**).

Thus, the presence of a visual landmark did not fundamentally alter the FEF transformation from an eye-centered visual code to an eye-centered gaze code. This preliminary result was important because it allowed us to narrow our focus to a more specific question: *given that the landmark clearly influences behavior (Fig. 1 B), how is this influence instantiated within the (primarily eye-centered) FEF visuomotor transformation? To* answer this, we asked if the influence of the landmark shift was somehow embedded in a more subtle way within those egocentric codes.

### Influence of the Landmark Shift on the Motor Code

We began our detailed allocentric analysis with a basic question: can the influence of the landmark shift be detected in the FEF motor responses? To test this, we performed an analysis similar to that shown above, but in this case by fitting RFs along the T-T’ continuum (Fig. 6). Fig. 6A-B shows visual and motor RF data from the same example neurons as Fig. 5, but now plotted in the T-T’ continuum. The visual RF is unchanged (Fig. 6A), as expected, since the landmark has not yet shifted. However, the best fit for the motor RF (Fig. 6B) had its best fit at the 3^rd^ step from T, indicated by the red arrowhead. In other words, it was shifted 30% toward T’ in landmark-fixed coordinates, i.e., similar to the observed behavior.

**Fig 6.**
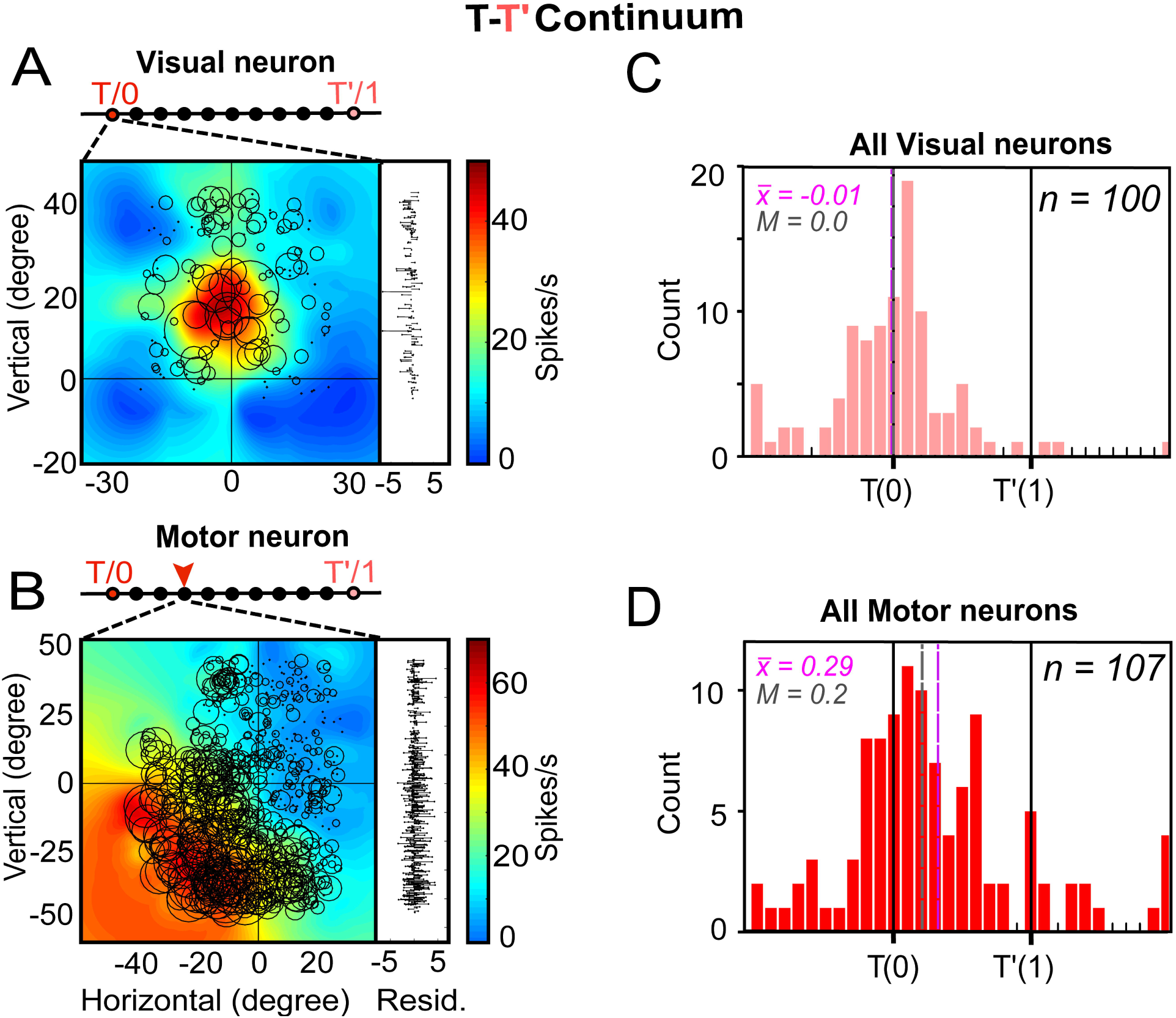
Allocentric influence on Visual and Motor responses. **(A-B)** Representations of visual and motor RFs using the same example neurons and graphic conventions as Figure 5, but here the fits were made along the T-T’ continuum. Notably, the best visual RF fit (a) was exactly on T (0), suggesting no influence of landmark on visual activity, whereas the best motor RF fit (b) was at the third step toward T’, suggesting a shift toward allocentric coordinates. **(C)** Frequency distribution of all spatially tuned visual neurons (n = 100) along the T-T’ continuum with the best fit at T (mean = – 0.01; median = 0) indicating no significant influence of T’ (p = 0.69, one sampled Wilcoxon signed rank test). **(D)** The distribution for spatially tuned motor neurons (n = 107) significantly shifted almost 1/3 from T toward T’ along the continuum (mean = 0.29; median = 0.2; p < 0.0001, one sampled Wilcoxon signed rank test).

These observations are documented for the entire population of neurons with significant spatial tuning in Fig 6C-D. These visual and motor populations overlapped with ∼ 95 % of the respective populations in Fig. 5, differing slightly because here we tested for tuning along the T-T’ continuum. Again, as expected, the visual response (Fig. 6C) peaked exactly at T with no significant shift toward T’ (mean = – 0.01; median = 0; p = 0.69, one sampled Wilcoxon signed rank test). However, the motor response (n = 107) displayed a significant shift (mean = 0.29; median = 0.2; p < 0.0001, one sampled Wilcoxon signed rank test) in the direction of the landmark shift (Fig. 6D), with a distribution significantly different from the visual response distribution (p = 0.0002; Mann-Whitney U-test). Notably, like the T-G transformation, the T-T’ shift was observed between the visual and the motor responses *within* visuomotor cells **(Supplementary Fig. S8).**

In summary, this analyses shows that the landmark shift influences the FEF motor code (Fig. 6D), but somehow this influence is embedded within the predominantly egocentric G code (Fig. 5F). Consistent with this, the motor fits produced significantly higher residuals for the T-T’ continuum than the T-G continuum (paired t-test, p < 0.0001), probably because the egocentric G model accounts for both the influence of the landmark shift and other internally-generated deviations from a perfect target code (T). To understand how this integration process occurs, we next examined when and how the allocentric influence becomes embedded into the egocentric code.

### Spatiotemporal analysis: when does the allocentric shift occur?

To track temporal transition of spatial coding T toward T’, we used a method developed previously for tracking the FEF T-G continuum during a memory delay (Sajad et al. 2016) (Fig. 7). Fig. 7A shows mean activity of our entire population of neurons, divided into 14 time-normalized bins proceeding from the onset of the visual response to the gaze shift onset (see Methods for details). Note that this period includes substantial spatially tuned delay activity that might be related to target memory and/or saccade planning. What might happen to spatial coding along the T-T’ continuum during this delay activity? As shown in Fig. 7B, the target code could immediately shift toward T’ (allocentric code, 1) after the landmark shift (after a brief visual delay), or this shift could be delayed until the final motor response (egocentric code, 2), or there might be a gradual transition (intermediate code, 3), as we observed previously for T-G.

**Fig. 7.**
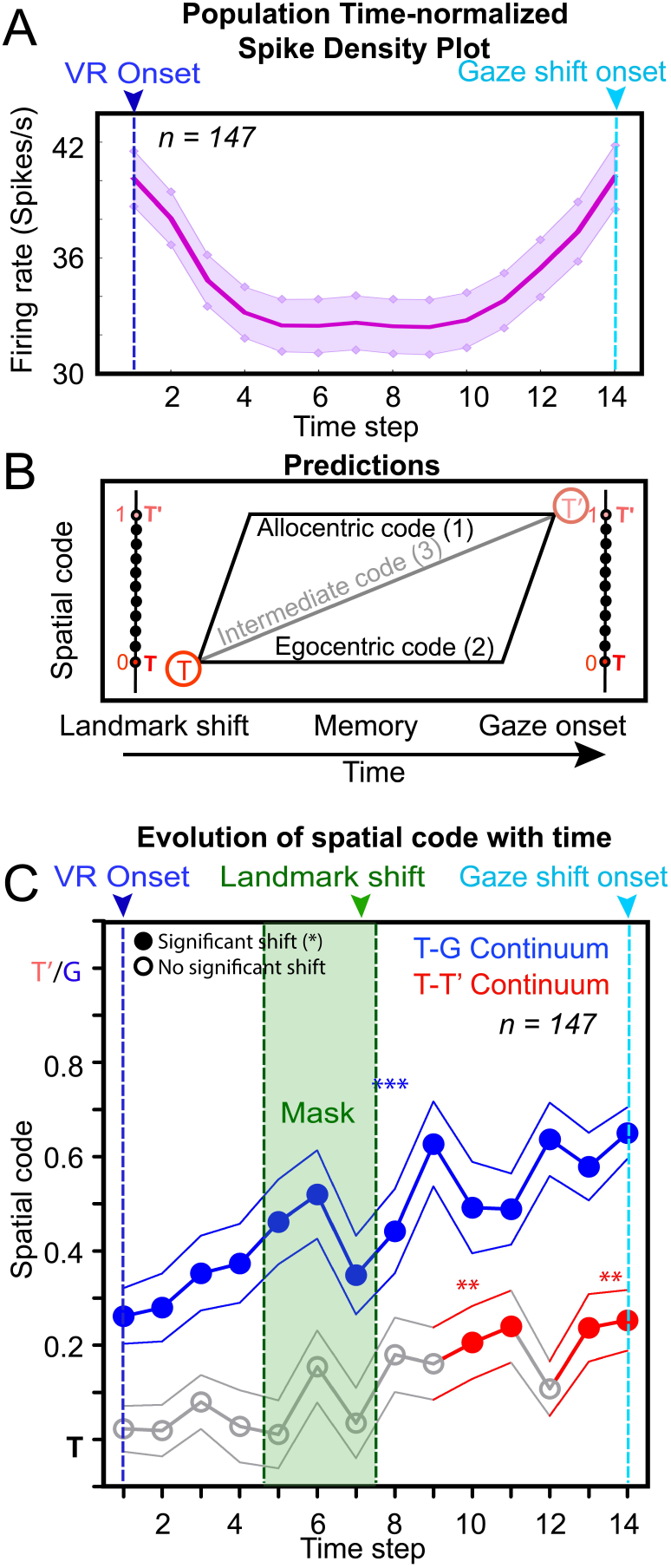
Spatiotemporal analysis at the populational level. **(A)** Time-normalized spike density plot (top 10 %) for whole of the population (n = 147) from visual response onset (VR onset, blue line) until the gaze shift onset (cyan line)**. (B)** Predictions for the transformation of code along the T-T’ continuum as the activity evolves from visual to motor through delay period: 1) target is transformed early on into an allocentric code and is then maintained in the memory for final execution/conversion 2) egocentric code, the target is maintained in the delay period and transformed later on, just before the movement initiation and 3) an intermediate code, a mix of 1 and 2, with a gradual transition from T to T’. **(C)** Progression of the spatial code (mean ± sem) along the T-G (blue curve) and T-T’ (red curve) continua with time (at different time steps) from VR onset until the gaze shift onset. Along the T-G continuum, the spatial coding starts at two steps from T and gradually builds up toward G with a significant shift toward G at each step. Note: along the T-T’ continuum, the activity starts to peak at 6^th^ step and gradually builds up toward T’ and gets significantly (p < 0.01, one sampled Wilcoxon signed rank test) embedded at steps 10 and 11 (i.e., three steps after Landmark shift) then the code reverts to T at step 12, finally shifting significantly at steps 13 and 14 (just before gaze shift onset). The shaded green area stands for the duration of mask and the green arrow corresponds to the landmark shift (L’); the solid blue and red circles represent a significant shift toward G and T’ respectively, with significant spatial coherence.

Fig. 7C shows the actual data: the mean (± sem) population fits for both the T-G continuum (blue data) and the T-T’ continuum (red data), plotted for each time step with respect to the major task events. (An example of this spatiotemporal analysis for a single visuomotor neuron is shown in **Supplementary Fig. S9).** Empty gray circles denote no significant shift from T, whereas colored circles represent significant shifts (p < 0.05, one sampled Wilcoxon signed rank test). As we observed previously (Sajad et al. 2015), the T-G fits immediately diverged from T after the visual response, following a series of intermediate codes toward G. In contrast, the T-T’ fits showed no significant shift until after the landmark shift at the end of the mask. But after that, importantly, the data showed two separate shift epochs: one transient shift about midway through the remaining delay period, and one later shift just before saccade motor burst (which, as we already saw, remains shifted). Overall, the pattern best resembled the ‘intermediate’ option illustrated in Fig. 7B, except for the slight dip that occurred at time step 12. However, this analysis does not yet account for the contribution of different cell types.

### Contribution of Different Cells and signals to the allocentric shift

To better understand what was happening during key period of landmark influence, we focused on a 7-step time-normalized interval between the landmark shift and saccade onset and broke the spatiotemporal analysis down into specific cell types (visual, visuomotor, and motor). The top row (from left to right) of Fig. 8 shows the overall activity of visual, visuomotor, and motor cells, including mean spike density plots for 1) all trials, 2) only the trials with the top 10% activity at each of time steps, and 3) and top 10 % activity from 80-180 ms after the landmark shift, i.e., when one might expect a visual response to the landmark shift. We then repeated our spatial analysis on each of these time steps. The bottom row of Fig. 8 again shows the T-G and T-T’ fits for these neuronal populations, using the same conventions as Fig. 7C.

**Fig. 8.**
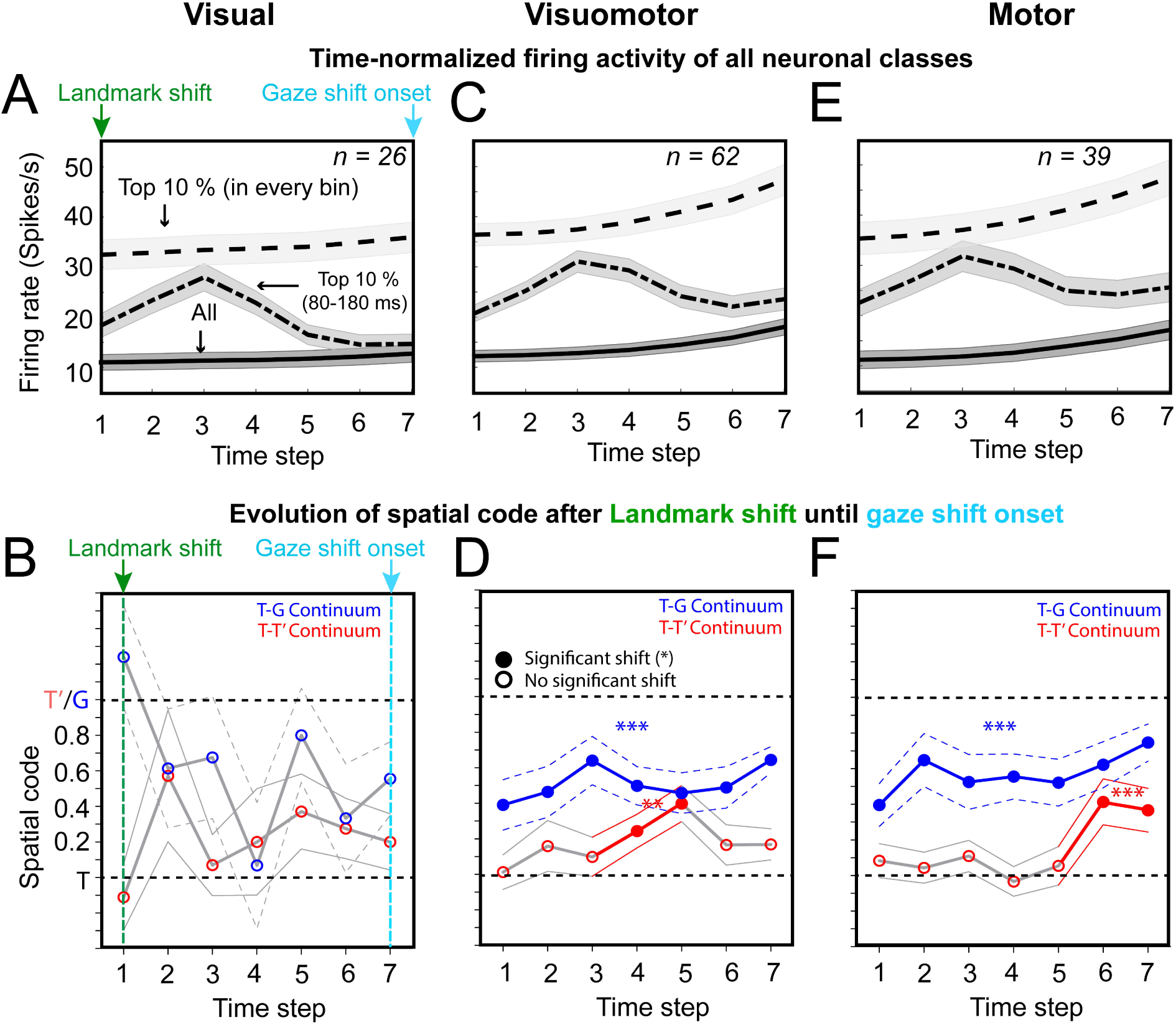
Spatiotemporal analysis of visual (V), visuomotor (VM) and motor (M) neurons aligned to landmark-shift (L’). **(A-B)** Spatiotemporal analysis of V neurons aligned to mask-off/landmark-shift (green arrow) until the gaze shift onset (cyan arrow). **(A)** Time-normalized (divided into 7 half-overlapping bins) spike density plot of V neurons (n = 26) for all trials (bottom trace), top 10 % trials in 80-180 ms window (middle trace) aligned to landmark-shift and top 10 % trials in each bin (top trace). **(B)** Progression of spatial code for V neurons from landmark-shift until the gaze shift onset. None of the steps (bins) was significantly shifted (p > 0.05, one sampled Wilcoxon signed rank test) toward G/T’ along both continua. **(C)** Time-normalized spike density plots for VM neurons (n = 62) as in **A**. **(D)** Progression of the spatial code for VM neurons along the T-G and T-T’ continua. Along the T-T’ continuum, there is a gradual shift of the code from the second step until it gets significantly shifted at steps 4 and 5, and shifts back at steps 6 and 7. Note: the opposite trend of the T-G and T-T’ curves — as the T-T’ activity starts to build up, the T-G curves advances in the opposite direction. **(E)** Time-normalized spike density plots for M neurons (n = 39)**. (F)** Progression of the spatial code for M neurons along the T-G and T-T’ continua. Notably, along the T-T’ continuum, the spatial code for M neurons remained constantly fixed around T for most of the steps before shifting toward T’ at steps 6 and 7 (just before gaze shift onset).

Several general observations can be gleaned from Fig. 8. First, although visual neurons (n = 26, Fig. 8A-B) retained some spatially tuned activity, they never showed a significant T-T’ shift (Fig. 8B; p > 0.05, one sampled Wilcoxon signed rank test), so we excluded them from further analysis (see methods for details). Second, in the remaining cell types (visuomotor and motor) the T-G (blue) and T-T’ (red) fits were not merely scaled versions of each other. The predictive power of neural activity for future gaze position (G) seems to peak at the 2^nd^ or 3^rd^ time step, whereas the influence of the landmark shift (T’) rises later. Further, in visuomotor cells these two continua appear to shift oppositely (toward each other) at the 4^th^ and 5^th^ step. Finally, any T-T’ shift that was observed in these cells occurred *after* the expected + 80-180 ms visual lag to the landmark (roughly at 3^rd^ time step), suggesting that additional processing time was required.

For visuomotor neurons (n = 62, Fig. 8C-D) there was a transient shift (Fig. 8D) in the target-related activity code toward T’ that starts building up at second step and significantly shifts at the fourth step (p = 0.02) and then peaking at the fifth step (shifts almost four spots toward T’; p < 0.05, One sampled Wilcoxon signed rank test) and then shifts back again toward T (the shift at steps 6 and 7 was not significant). In short, visuomotor cells were responsible for the transient shift in target coding that occurred during the second memory interval after the physical landmark shift.

Motor neurons (n = 39) showed an opposite trend to the VM neurons (n = 39, Fig. 8E-F); the spatial codes along the T-T’ continuum remained fixed around T for most of the steps and a significant shift (p < 0.05, One sampled Wilcoxon signed rank test) toward T’ was only during the final steps (6 and 7) for gaze command. Notably, the spatial-code evolution profile of VM neurons at fourth and fifth step was significantly different for the corresponding step of M neurons (p < 0.05, unpaired t-test), implying that their encoding trajectories are different/opposite (this is also evident from T-T’ curve in Figure 5 — the first shift is possibly attributed to visuomotor neurons whereas the latter shift maybe related to motor neurons). In summary, shortly after the T-T’ shift disappeared in visuomotor neurons, it reappeared in motor neurons, lasting until the saccade burst.

### Integration of allocentric shift in the egocentric motor code

Thus far, we have observed that the landmark shift induces a shift in T-T’ coordinates at various times in visuomotor and motor neuron activity, but we still need to reconcile this with the fact that G (in eye coordinates) dominates the motor code, and yet somehow incorporates the T-T’ shift. To address this point, we re-plotted the T-T’ as a function of the corresponding T-G for each neuron that was spatially tuned to both (Fig. 9A). We hypothesized that this allocentric/landmark influence could be encoded in the FEF neural activity in four possible ways (Fig. 9B): 1) No influence (which we already showed is not the case), 2) an independent influence (in which case there would be no relationship between T-G and T-T’), 3) an integrated influence (in which case T’ would vary from T as a function of G), or 4) partial integration, i.e., some intermediate between the latter two options. In the latter two cases this would directly implicate the T-T’ fits in behavior, since G is derived directly from behavioral measures and the T-G continuum indicates the degree to which neural activity influences gaze as opposed to coding target location.

**Fig. 9.**
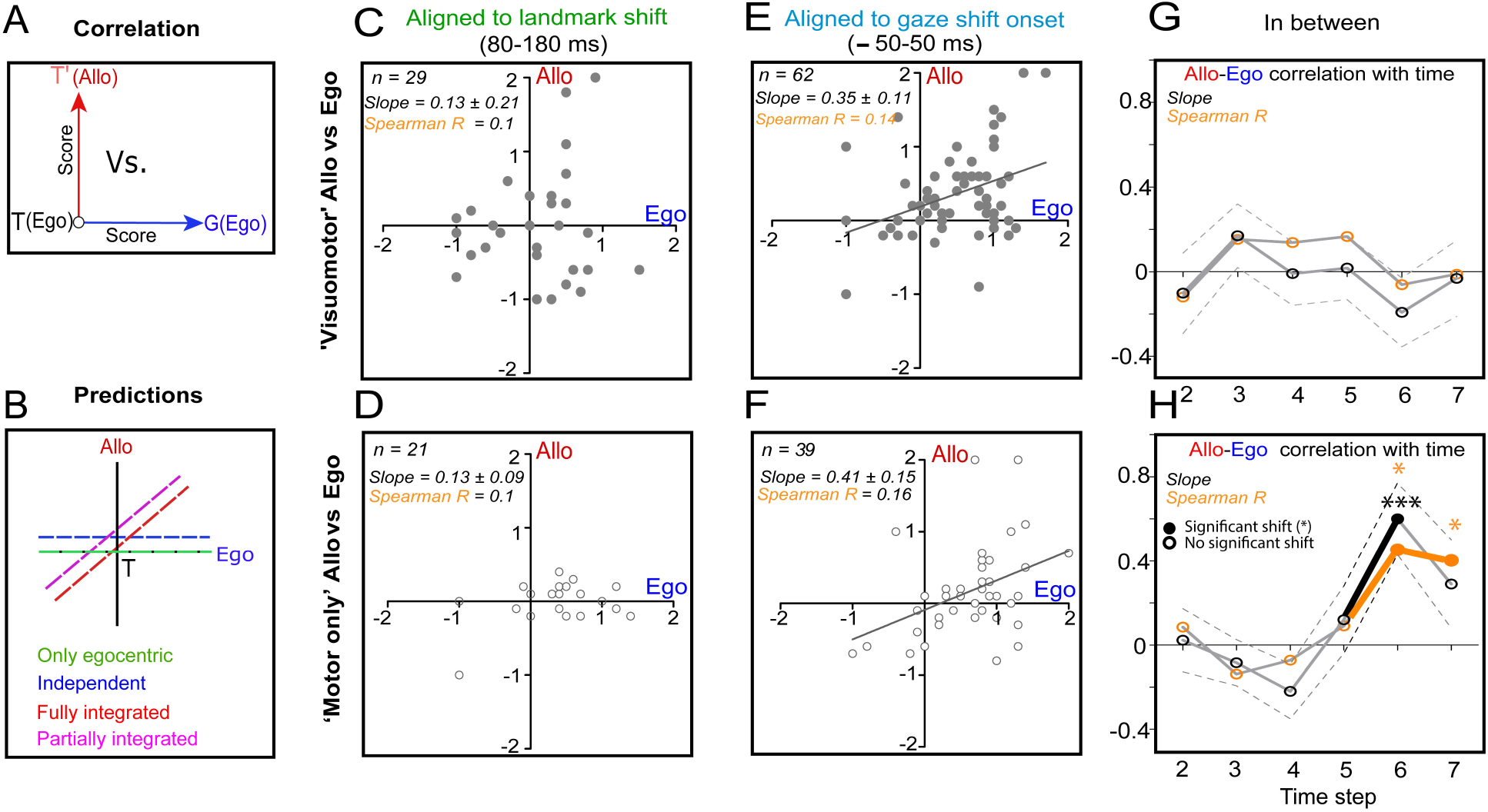
Correlation between T-G (Ego-Ego, x-axis) and T-T’ (Ego-Allo, y-axis) scores and the spatiotemporal evolution of Allo-Ego correlation. **(A)** A schematic of the correlation computation between T-T’ and T-G scores (best point along the continuum). Different predictions for the embedding of allocentric (T-T’) shift in the egocentric (T-G) code: 1) only egocentric, or 2) independent encoding, or 3) fully integrated, or 4) partially integrated. **(C)** No significant correlation between the corresponding T-T’ and T-G scores of VM neurons in the 80-180 ms window aligned to landmark shift (n = 29, Spearman R = 0.01, slope = 0.13 ± 0.29, p = 0.55). **(D)** No significant correlation between the corresponding T-T’ and T-G scores of M neurons in the 80-180 ms window aligned to landmark shift (n = 21, Spearman R = - 0.05, slope = 0.13 ± 0.09, p = 0.20). **(E)** Significant correlation between the corresponding T-T’ and T-G scores of VM neurons in the perisaccadic window (n = 62; Spearman R = 0.31; slope = 0.35 ± 0.11, Intercept = 0.19 ± 0.09, p < 0.05). **(F)** Significant correlation between the corresponding T-T’ and T-G scores of M neurons in the perisaccadic window (n = 39; Spearman R = 0.39; slope = 0.41 ± 0.15, Intercept = - 0.09 ± 0.13, p < 0.05). There was a significant difference between the intercepts of M and VM neurons in the perisaccadic window (F = 4.54, p = 0.036). **(G)** No significant correlation between T-T’ and T-G scores in the VM neurons with time (from steps 2 until 7, corresponding to Figure 6). Note: Steps 2 to 7 included the time between the transient 80-180 ms response aligned to L’ and the – 50-50 ms perisaccadic burst. **(H)** A significant correlation developed between T-T’ and T-G scores of M neurons at steps 6 (Spearman R = 0.45, p < 0.05; Slope = 0.60 ± 0.17) and 7 (Spearman R = 0.40, p < 0.05; Slope = 0.29 ± 0.21) with time.

First, we performed this test on the period (80-180 ms) aligned to the landmark shift, when one would expect the first visual influence (Fig. 9C-D). We found no significant correlation between the T-T’ and T-G scores for either visuomotor (Spearman R = 0.01, slope = 0.13 ± 0.29; p = 0.55) or motor cells (Spearman R = −0.05, slope = 0.13 ± 0.09; p = 0.20) neurons. However, when we repeated this test on the peri-saccadic motor response (Fig. 9E-F), we found significant correlations for both (Visuomotor: Spearman R = 0.31, slope = 0.35 ± 0.11; Intercept = 0.19 ± 0.09, p < 0.05; Motor: Spearman R = 0.39; slope = 0.41 ± 0.15, Intercept = - 0.09 ± 0.13, p < 0.05). Although the visuomotor and motor population slopes were not significantly different from each other (F = 0.10; p = 0.75), their intercepts were significantly different (F = 4.54; p = 0.036), suggesting further differences between the contributions of these cell types. The combined analysis for motor neurons (M + VM, n = 101) also showed a significant correlation between T-T’ and T-G scores (Spearman R = 0.31; slope = 0.35 ± 0.09, Intercept = 0.09 ± 0.08, p < 0.05). Thus, these egocentric and allocentric signals became at least partially integrated, sometime before or at the motor burst.

To track when this integration occurred, we computed the T-G/T-T’ spearman correlations and slopes (Fig. 9G-H) for the time epochs (from steps 2-7 that included the time between the 80-180 ms transient response to L’ and the -50-50 ms perisaccadic burst) used in the preceding figure. We found no significant correlation (p > 0.05) in visuomotor neurons (Fig. 9G), whereas a significant correlation (at step 6 and 7) developed in motor neurons, i.e., as the saccade command was imminent. This analysis shows that the initial transient T-T’ visuomotor shift shown in Fig. 8D is independent of the egocentric target memory code, whereas the integration of these signals only arises in late motor neuron response, somehow spreading/integrating then across all motor responses during the saccade. This also shows that cells with codes that shift toward T’ also tend to shift more toward a gaze motor code (T toward G). Since G is based on actual behavioral measurements, this means that these cells tend to influence the behavioral shift more than ‘non-shifters’.

## DISCUSSION

In the current study, we combined a cue-conflict gaze task, in which a visual landmark shifted in the interval between seeing and acting on the object, with a model-fitting analysis approach to examine if, when, and how the FEF contributes to the integration of egocentric and allocentric visual cues for aiming gaze direction. As noted above, an advantage of our model-fitting method is that it not only shows the presence of specific types of information (e.g., T, G), it also allows us to test hypotheses about how this information is coded and multiplexed, for example in different frame of reference (e.g. relative to the eye vs. landmark). In summary, we found that: 1) In the presence of a visual landmark, FEF visual and motor responses continued to be dominated by egocentric codes, i.e. T (Target-relative to eye) and G (future gaze relative to eye), but embedded within the motor code was a partial shift toward landmark-centered coordinates (T’). 2) The timing of this shift was cell-type dependent: it was observed first as a transient unintegrated shift (T-G and T-T’ not correlated) in the visuomotor target memory response, and later as an integrated shift (T-G and T-T’ correlated) in the build-up response of motor neurons. 3) Finally, an integrated shift was observed in both visuomotor and motor neuron responses during the actual gaze shift.

### 1. Relationship to previous human studies

Previous human neuropsychological and neuroimaging studies have suggested that egocentric and allocentric information get separated early on in the visual areas and are associated with ‘dorsal stream’ parietal cortex, and ‘ventral stream’ temporal cortex respectively (Goodale and Haffenden 1998; Byrne and Crawford 2010; Chen et al. 2014; Filimon 2015) and then are recombined in the fronto-parietal cortex for behavior (Chen, Monaco, et al. 2018). Our current results clearly support the latter notion, providing a specific mechanism for allocentric-egocentric integration of gaze in the frontal eye fields. It has also been argued that all of the brain’s spatial codes, even the allocentric (Filimon 2015), are ultimately egocentric because they derive from egocentric sensory inputs and control egocentric motor commands. Again, our data support this notion, in the sense that our FEF Visual/Motor codes were predominantly egocentric (Te, Ge), and allocentric information was progressively integrated into these codes. Our data cannot comment directly on earlier visual codes, but there does appear to be a special need for keeping track of relative position for allocentric coding, likely the reason why this involves ventral stream vision (Milner and Goodale 2006; Thaler and Goodale 2011). Further, our time-course data are qualitatively consistent with the observations from neuropsychological studies of dorsal vs. ventral stream damage that allocentric codes lag behind egocentric codes when used to guide action (Goodale and Haffenden 1998; Khan et al. 2005; Milner and Goodale 2006).

### 2. Relationship to previous neurophysiology studies

Until now, what we know about integration of egocentric and allocentric visual cues for action comes from human psychophysical studies. Previous experiments (Colby 1998; Lemay et al. 2004; Neggers et al. 2005; Mou et al. 2006; Neely et al. 2008; Byrne and Crawford 2010; Thaler and Goodale 2011; Fiehler et al. 2014; Klinghammer et al. 2017b; Li et al. 2017) suggest that egocentric and allocentric cues are optimally combined at the behavioral level for reaching and saccades and thus several theories have been postulated for this optimal integration (Körding and Wolpert 2004; Byrne et al. 2007; Körding et al. 2007; Byrne and Crawford 2010; Filimon 2015; Klinghammer et al. 2017b). Most of these theories rely on Bayesian concepts of statistical integration (Körding and Wolpert 2004; Körding et al. 2007; Mutluturk and Boduroglu 2014; Aagten-Murphy and Bays 2019). In this framework, the brain represents the prior statistical distribution of the target/task and the level of uncertainly in the sensory feedback, thus potentiating the combining of both kinds of information and allowing performance-optimization. Byrne and Crawford (Byrne and Crawford 2010) were the first to propose this, using a maximal likelihood estimator (MLE) model, where the amount of reliable information from both cues determines the relative/optimal combining of both cues. The current results, both behavioral and neural, qualitatively and quantitatively agree with those studies, but cannot formally claim optimal integration because this would require explicitly training monkeys to do pure egocentric and pure allocentric controls. We did not wish to do this because it could be argued that this might influence or even create our results.

Several animal studies have sought to distinguish allocentric versus egocentric codes in auditory cortex (Town et al. 2017), in parietal cortex (Chen, DeAngelis, et al. 2018), and in hippocampus during navigation (Matsumura et al. 1999; Rolls 1999). But to our knowledge, very few neurophysiological studies (Olson and Gettner 1995; Dean and Platt 2006; Uchimura, M; Kumano, H; Kitazawa 2017) have focused on distinguishing egocentric and allocentric codes in visual and visuomotor systems. A saccade study reported that a population of posterior cingulate cortex (CGp) neurons encodes visuospatial events in allocentric coordinates (Dean and Platt 2006), whereas another study showed that only a fraction (∼10%) of precuneus neurons encoded the allocentric location of a dot in the presence of a background landmark (Uchimura, M; Kumano, H; Kitazawa 2017). Experiments, wherein monkeys had to saccade to a target relative to another cue in space have shown that some neurons in the SEF (Olson and Gettner 1995; Tremblay et al. 2002) and lateral intraparietal cortex (Sabes et al. 2002) can encode object-centered information, i.e. one particular part of an object. However, none of these studies examined the question of how landmark-centered and egocentric visual cues are integrated and weighted at the cellular level for motor behavior.

### 3. Neurophysiological Mechanisms of Ego-Allocentric Integration for Gaze

Based on our results, we propose the circuit model shown in Fig. 10. We will now explain this model while citing the relevant evidence from our study and the previous literature. First, based on previous studies discussed above (Byrne and Crawford 2010; Chen et al. 2014; Filimon 2015), we assume that there is at least a relative segregation of egocentric (blue arrow, dorsal) and allocentric information (red arrow, ventral) early in the visual system. The ‘dorsal stream’ egocentric system would provide the default drive for the FEF, providing low-latency inputs in target-relative-to-eye coordinates (Fig. 5C **and** Fig. 6C). We speculate that the SC and the SEF that share reverse connections with the FEF are also involved in this integration process, however, the SC (that receives connections from FEF for gaze output command) may integrate these signals more optimally for final behavior.

**Fig. 10.**
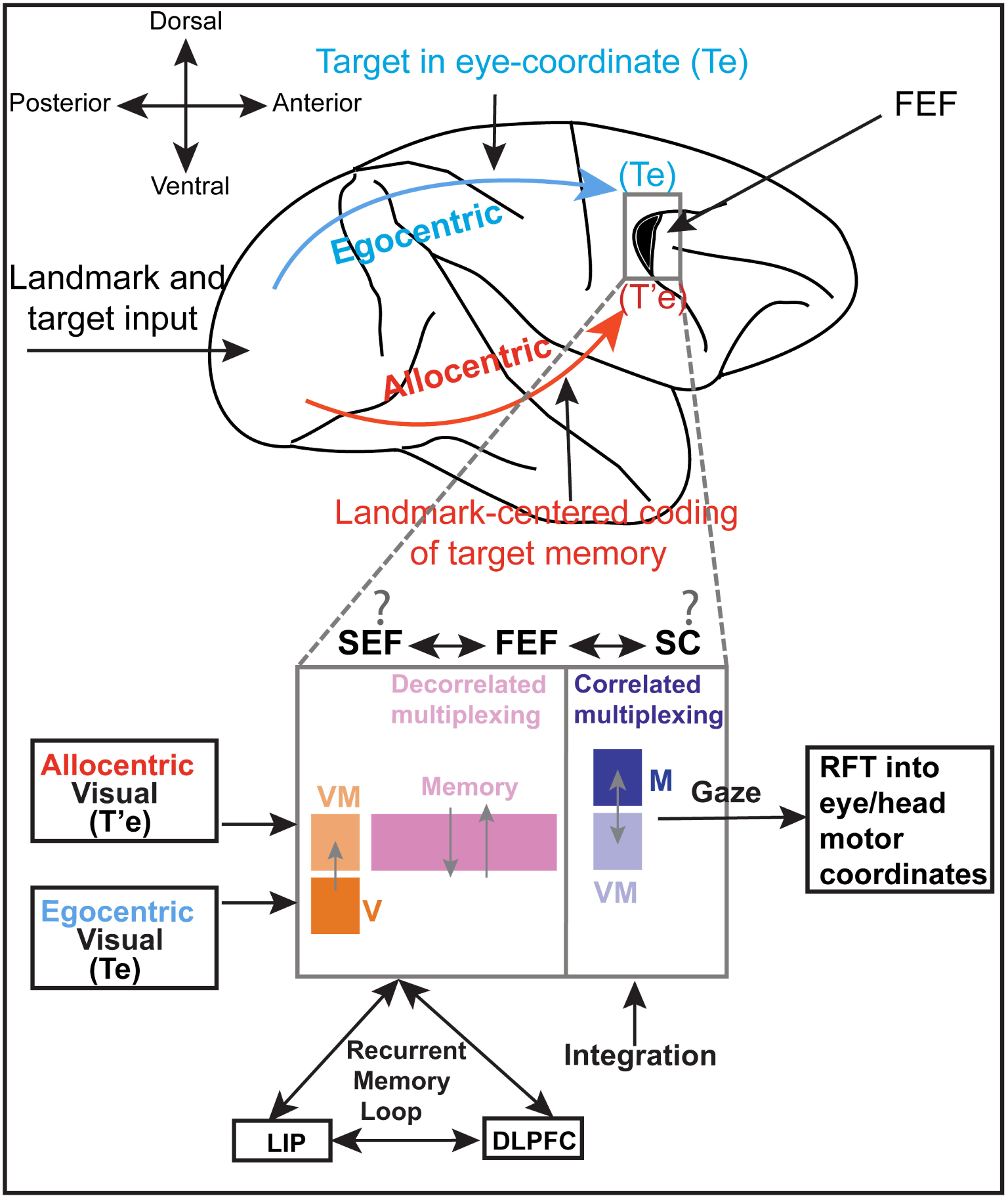
A neural circuit model for egocentric and allocentric multiplexing of signals. As the information (target and landmark) enters the early visual areas, it gets separated into two pathways (top): the egocentric information (blue arrow) takes the dorsal stream with the target in eye(Te), and the allocentric information (red arrow) follows the ventral stream, wherein the landmark-centered memory is updated that finally converges in the FEF (gray box, and recurrent connections with SC and SEF) with the egocentric flow. Bottom: A circuit diagram for integration of egocentric and allocentric codes in FEF (zoomed-in gray box). The visual information from extrastriate areas enters the FEF circuit as two different inputs — an egocentric (Te) input to the visual (V, dark orange) neurons and an allocentric input to visuomotor neurons (VM, light orange). This sensory information is then gated to appropriate memory network (purple, primarily VM neurons), wherein the decorrelated scaling/multiplexing occurs between the egocentric and allocentric flows. The VM neurons primarily reflect more processed visual information possibly due to gating in the fronto-striato-thalamic loop (Chatham and Badre 2015). Finally, as the gaze is imminent, the VM neurons (light blue) pass on the information of most recent multiplexed memory to the M neurons (dark blue), while getting fully integrated in the final motor burst (gaze command) through transient activation of VM and M synapses and thus potentiating a reference frame transformation (RFT) in eye/head coordinates. Note: the multiplexing of egocentric and allocentric signals may get continuously updated through a working memory loop between the LIP, DLPFC and FEF. The gray arrows indicate the flow between neuronal classes.

Because the FEF participates in a distributed working memory loop that involves lateral intraparietal cortex (LIP), dorsolateral prefrontal cortex (DLPFC) and other brain areas (Christophel et al. 2017; Pinotsis et al. 2019), we now propose that at some point at or before FEF, ventral stream allocentric signals are incorporated, influencing target memory at the level of FEF visuomotor neurons after a slight delay (Fig. 8D). Further, we suggest that similar working memory mechanisms may be central to integration of egocentric and allocentric information in LIP and DLPFC. The observed amplitude and independent nature of this signal (Fig. 9G) supports the notion that this signal is already pre-weighted (presumably based on past experience and various perceptual rules), and arises independently, not yet being fully integrated even at this late stage in the system. It appears that this signal dissipates rapidly (at least in our task), and yet still must be stored somehow, either in a distributed activity fashion through the memory loop (Christophel et al. 2017; Miller et al. 2018; Pinotsis et al. 2019) or as temporary synaptic changes (Mayford et al. 2012; Masse et al. 2019). It then reemerges in the motor neuron build-up response (Fig. 8E), but this time in a more integrated form (Fig. 9H). Thus, the visuomotor-to-motor circuitry seems to be the key integration point, but this could also involve signal shuffling across the distributed, reentrant memory network (Christophel et al. 2017; Pinotsis et al. 2019). Indeed, there seems to be some spatio-temporal push-pull between these two neuronal classes when comparing their codes through time in our task (Fig. 8). This integration of egocentric and allocentric cues is reminiscent of ‘bump attractor’ simulations wherein two memory ‘bumps’ influence each other and finally turn into one bump (Wimmer et al. 2014). Finally, presumably through shared interconnections, both visuomotor and motor cells arrive at a similarly integrated code with oppositely cancelling biases (Fig. 9E-F), and then providing this to the downstream motor system (i.e., SC, reticular formation, etc.).

### 3. General Implications

An important aspect of our task is that we did not explicitly train animals to follow an egocentric or allocentric strategy, but instead rewarded them for either strategy, or some intermediate combination. We did this to avoid altering the normal circuitry, and also in hopes that the results would generalize well to real-world circumstances. The fact that the behavioral results resemble human reach and saccade results supports this notion and suggests our results would generalize well to other species and systems. In one respect our task was not realistic: to test between the influence of visual landmarks, it was necessary to introduce a cue conflict in a delay task. But based on our previous results it appears that such signal transformations scale down in time. We would thus expect our model to generalize to normal saccade delays in the presence of a stable landmark. Although the specific mechanisms used by the FEF to integrate egocentric and allocentric information appear to be novel, the general principle of multiplexing spatial information within and across neurons is common in visual and memory systems (Beck et al. 2008; Blohm and Crawford 2009; Campos and Segraves 2011; Meister et al. 2013; Ratté et al. 2013). Finally, it is noteworthy that egocentric versus allocentric functions appear to be differentially preserved after large-scale damage to the dorsal versus ventral visual stream (Milner and Goodale 2006), although more subtle integration problems likely have not yet been explored. The promise here is that the fundamental knowledge of these semi-redundant mechanisms, in conjunction with new developments in brain plasticity, might benefit rehabilitation of patients with brain damage and disease.

## CONTRIBUTION

VB did the experiments, analyzed the data and wrote the manuscript. AS contributed to data analysis. XY helped in the technical aspects of the recording data. JL trained the animals. HW performed the surgeries and helped in neural recordings. JDC conceived the study and contributed to writing and editing of the manuscript.

## ACKNOWLEDGEMENT

This project was supported by a Canadian Institutes for Health Research (CIHR) Grant and the Vision: Science to Applications (VISTA) Program, which is supported in part by the Canada first Research Excellence Fund. VB, XY, and HW were supported by CIHR and VISTA. AS was supported by an Ontario Graduate Scholarship. JDC is supported by the Canada Research Chair Program.

