## Supplementary file for "Integration of eye-centered and landmark-centered codes in frontal eye field gaze responses"

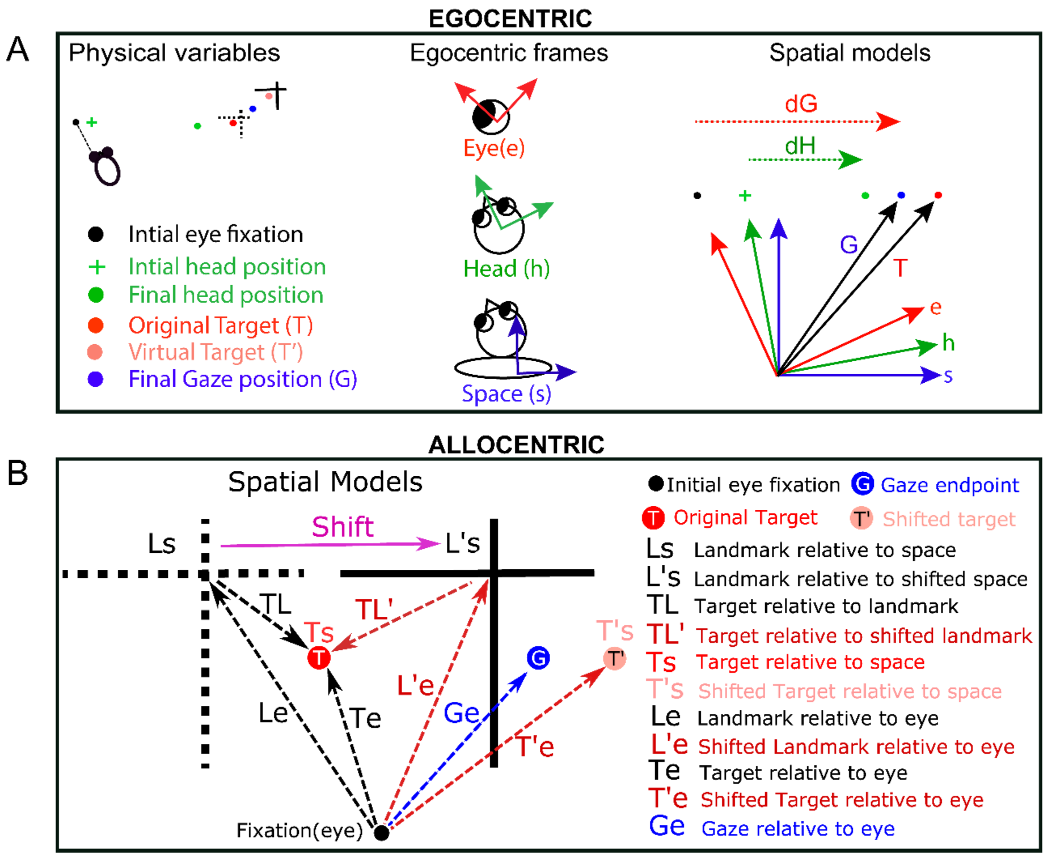


**Supplementary Fig. S1: List of all the egocentric and allocentric models tested (A)** Different egocentric models that were tested. Left: Different physical variables for one trial. Middle: three basic egocentric frames (eye, head and body/space). Different canonical models obtained after plotting different physical parameters in basic egocentric frames. dG: displacement of gaze; i.e., final gaze position with respect to the fixation point (not the eye). dH: head displacement in space coordinates. dE (not shown) is eye displacement in the head that accompanies the gaze shift. **(B)** Different allocentric models (see below) that were tested along with the most relevant egocentric models (Te, Ge and Ts). The broken black cross stands for the initial landmark (L) location whereas the solid black cross corresponds to the shifted (indicated by pink arrow) landmark (L'). The red, the blue and the light red solid circles represent the target (T), the gaze endpoint (G) and the virtually shifted target (T') locations respectively. Allocentric models tested: Ls, landmark relative to space; Le, landmark relative to eye; TL, target relative to landmark; L's, shifted landmark relative to space; L'e , shifted landmark relative to eye, TL', target relative to shifted landmark, T'e, virtually shifted target relative to eye; T's virtually shifted target relative to space.

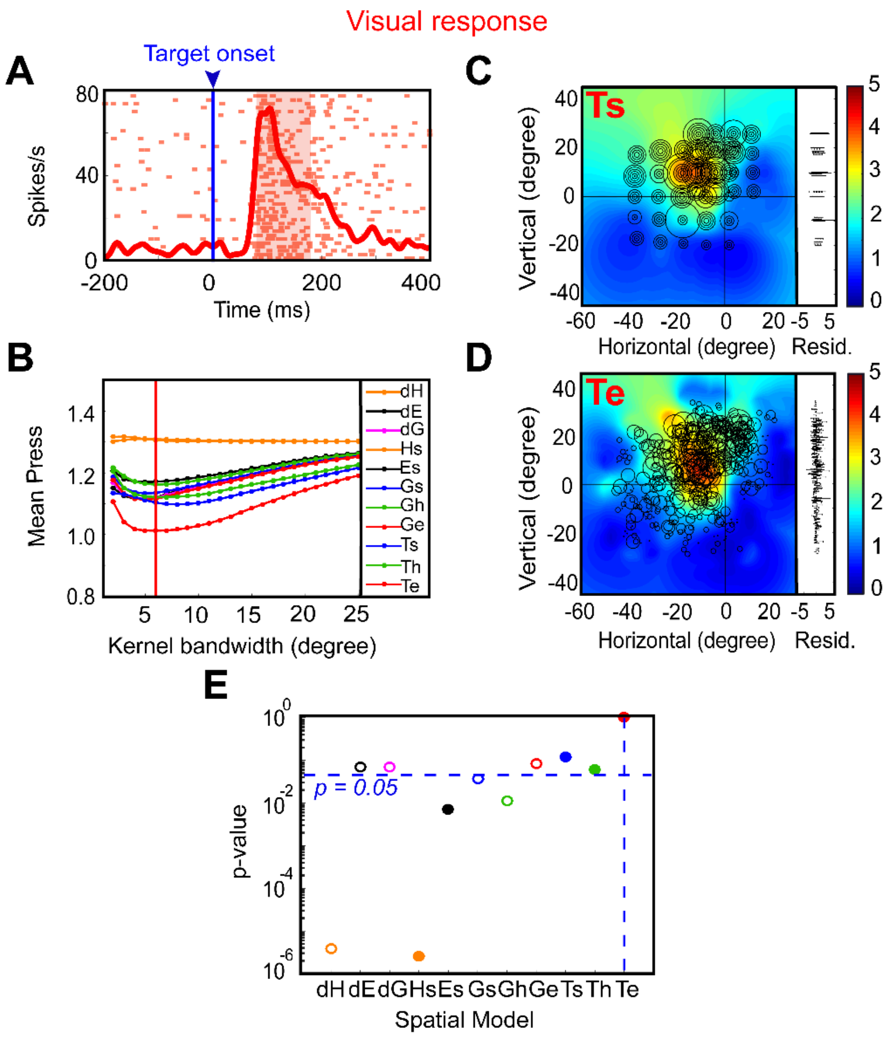


**Supplementary Fig. S2: An example of analysis of a visual response. (A)** Raster and spike density plot (red curve) of a visually responsive neuron aligned to target on (blue line). The red-shaded area corresponds to the sampling window for spatial-model dissociation analysis. **(B)** The mean residuals from the PRESS-statistics for all tested models as a function of different kernel bandwidths (2-25°). The red vertical line stands for the lowest mean PRESS for Te (target relative to eye) with a kernel bandwidth of 6°. **(C-D)** Representation of activity in two spatial models: Ts: target in space (screen) and Te: target relative to the eye. The circle represents the magnitude of the response and heat map shows the non-parametric fit to these data (red blob highlights the hot spot of the RF of neuron). The corresponding residuals are shown to the right. **(E)** The p-values for statistical analysis and comparison between Te ($p = 10^0 = 1$; the best-fit model with lowest residuals) and other models (Brown-Forsythe test).

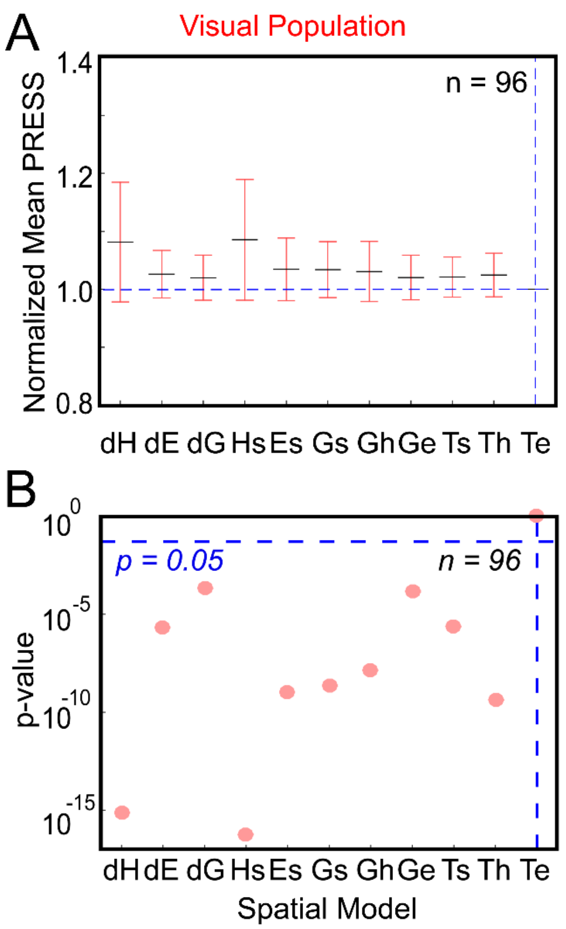


**Supplementary Fig. S3: Egocentric population fit residuals and statistics for all visual neurons. (A)** The population mean (± SEM) across the mean PRESS residuals of all visually responsive neurons (n = 96), i.e., data relative to fits made to all tested egocentric models. These values have been normalized by dividing by the mean PRESS residuals of the best fit model, in this case Te. **(B)** P-Value statistics done on the residuals shown above (Brown-Forsythe test). Te (broken vertical blue line) is the best fit and all other models were eliminated, i.e. $P < 0.05$.

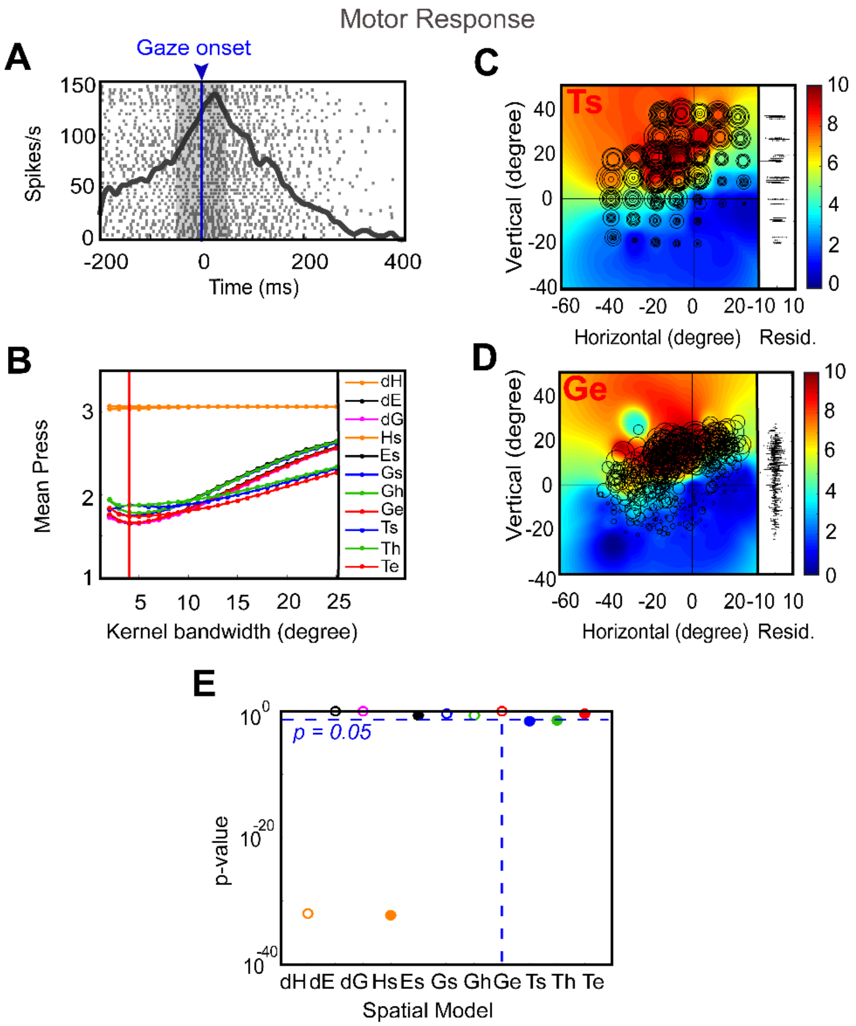


**Supplementary Fig. S4: An example of analysis of a motor response. (A)** Raster and spike density plot (red curve) of a gaze responsive (movement) neuron aligned to gaze on (blue line). The gray-shaded area corresponds to the sampling window for spatial-model dissociation analysis. **(B)** The mean residuals from the PRESS-statistics for all tested models as a function of different kernel bandwidths (2-25°). The red vertical line stands for the lowest mean PRESS for Ge (gaze relative to eye) with a kernel bandwidth of 4°. **(C-D)** Representation of activity in two spatial models: Ts: target in space (screen) and Ge: gaze relative to the eye. The circle represents the magnitude of the response and heat map shows the non-parametric fit to these data (red zone represents the hot spot of the RF of neuron). The corresponding residuals are shown to the right. **(E)** The p-values for statistical analysis and comparison between Ge ($p = 10^0 = 1$; the best-fit model with lowest residuals) and other models (Brown-Forsythe test).

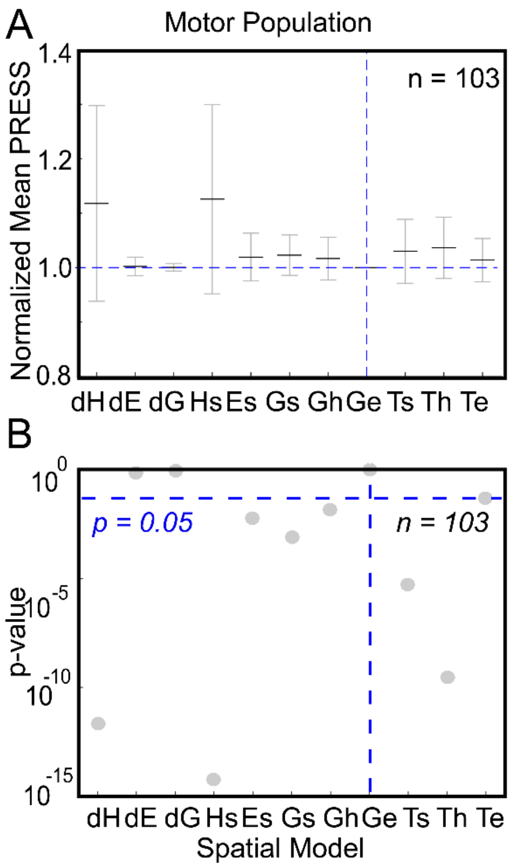


**Supplementary Fig. S5: Egocentric population fit residuals and statistics for all motor neurons. (A)** The population mean (± SEM) across the mean PRESS residuals of all motor neurons (n = 103), i.e., data relative to fits made to all tested egocentric models. These values have been normalized by dividing by the mean PRESS residuals of the best fit model, in this case Ge. **(B)** P-Value statistics done on the residuals shown above (Brown-Forsythe test). Ge (broken vertical blue line) is the best fit (however, dE and dG were not eliminated) and all other models were eliminated, i.e. $P < 0.05$.

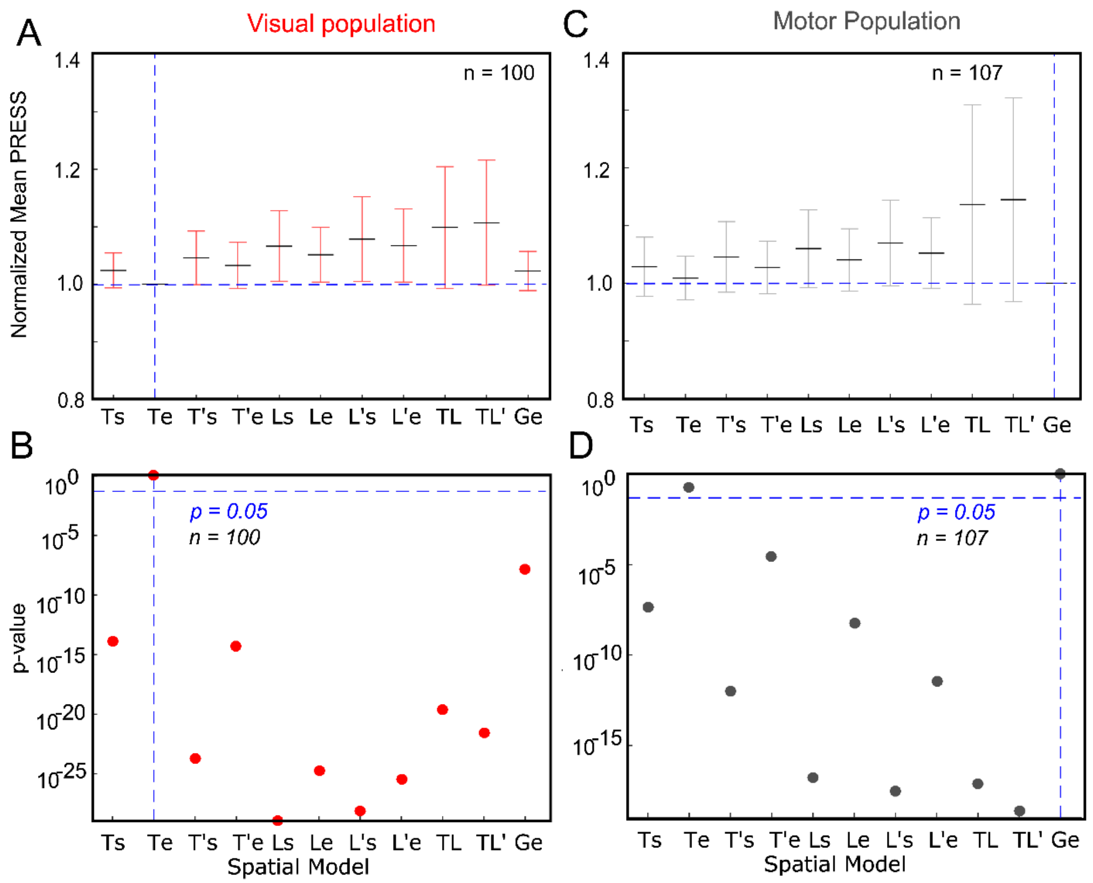


**Supplementary Fig. S6. Allocentric population fit residuals and statistics for all visual and motor neurons. (A)** The population mean (± SEM) across the mean PRESS residuals of all visually responsive neurons (n = 100), i.e., data relative to fits made to all tested models. These values have been normalized by dividing by the mean PRESS residuals of the best fit model, in this case Te. **(B)** P-Value statistics done on the residuals shown above (Brown-Forsythe test). Te (broken vertical blue line) is the best fit and all other models were eliminated, i.e. $P < 0.05$. **(C)** The population mean (± SEM) across the mean PRESS residuals of all motor neurons (n = 107), i.e., data relative to fits made to all tested models. These values have been normalized by dividing by the mean PRESS residuals of the best fit model, in this case Ge. **(D)** P-Value statistics done on the residuals shown above (Brown-Forsythe test). Ge (broken vertical blue line) is the best fit (however, Te is not eliminated) and all other models were eliminated, i.e. $P < 0.05$

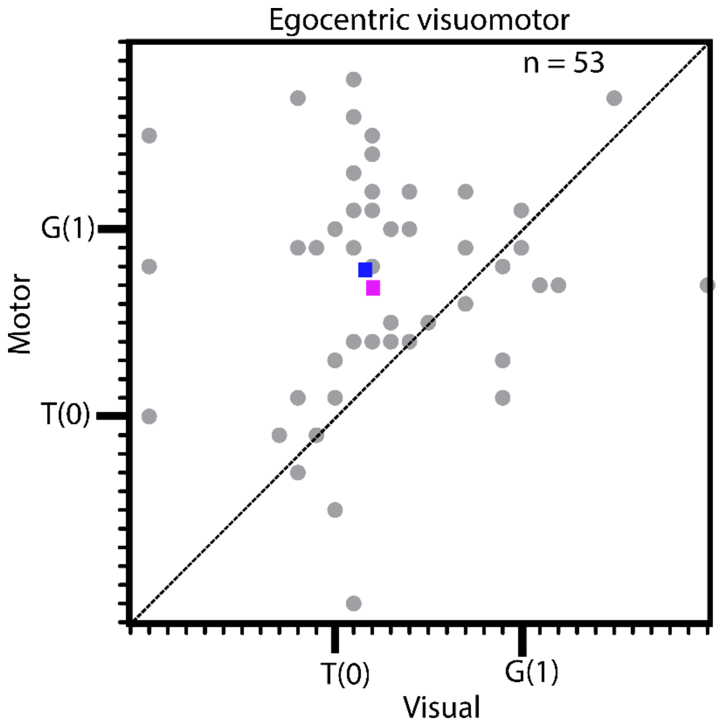


**Supplementary Fig. S7: Visuomotor transformation at the single cell (visuomotor neurons) along the T-G continuum.** Movement response (y-axis) as a function of the corresponding visual response (x-axis) for VM neurons. The gray dot represents visual best-fit model at x and the movement best-fit model at y. The blue and the pink squares represent the median and the mean, respectively, of the best-fit points for the movement against the best-fit point of the visual response. There was a significant shift ($p < 0.0001$, Wilcoxon matched-pairs signed rank test) toward G (most dots shifted above the broken unity line toward G suggesting a transformation from the target (T) to gaze (G) coding.

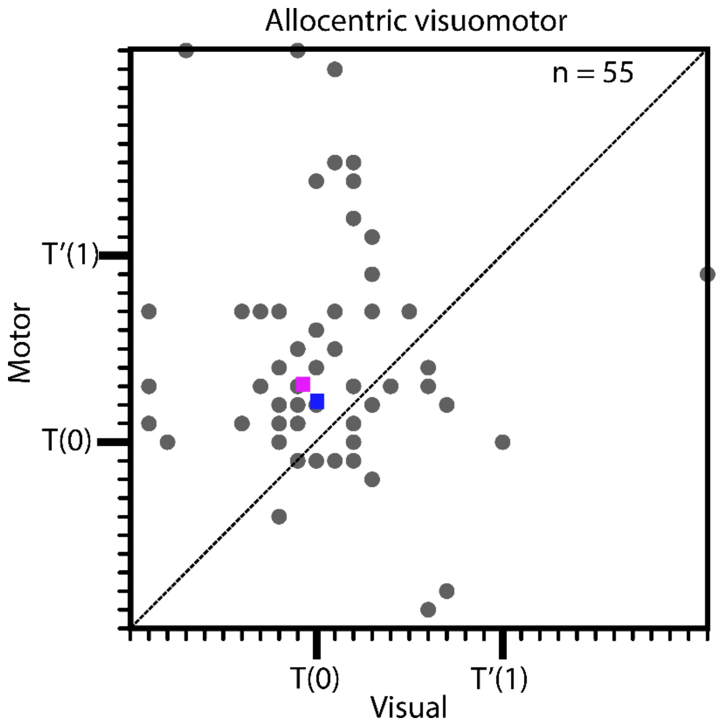


**Supplementary Fig. S8: Visuomotor transformation at the single cell (visuomotor neurons) along the T-T’ continuum.** Movement response (y-axis) as a function of the corresponding visual response (x-axis) for VM neurons. The gray dot represents visual best-fit model at x and the movement best-fit model at y. The blue and the pink squares represent the median and the mean, respectively, of the best-fit points for the movement against the best-fit point of the visual response. There was a significant shift ($p = 0.0006$, Wilcoxon matched-pairs signed rank test) toward T’ (most dots shifted above the broken unity line toward T’) suggesting a transformation from the target (T) to shifted target (T’) coding.

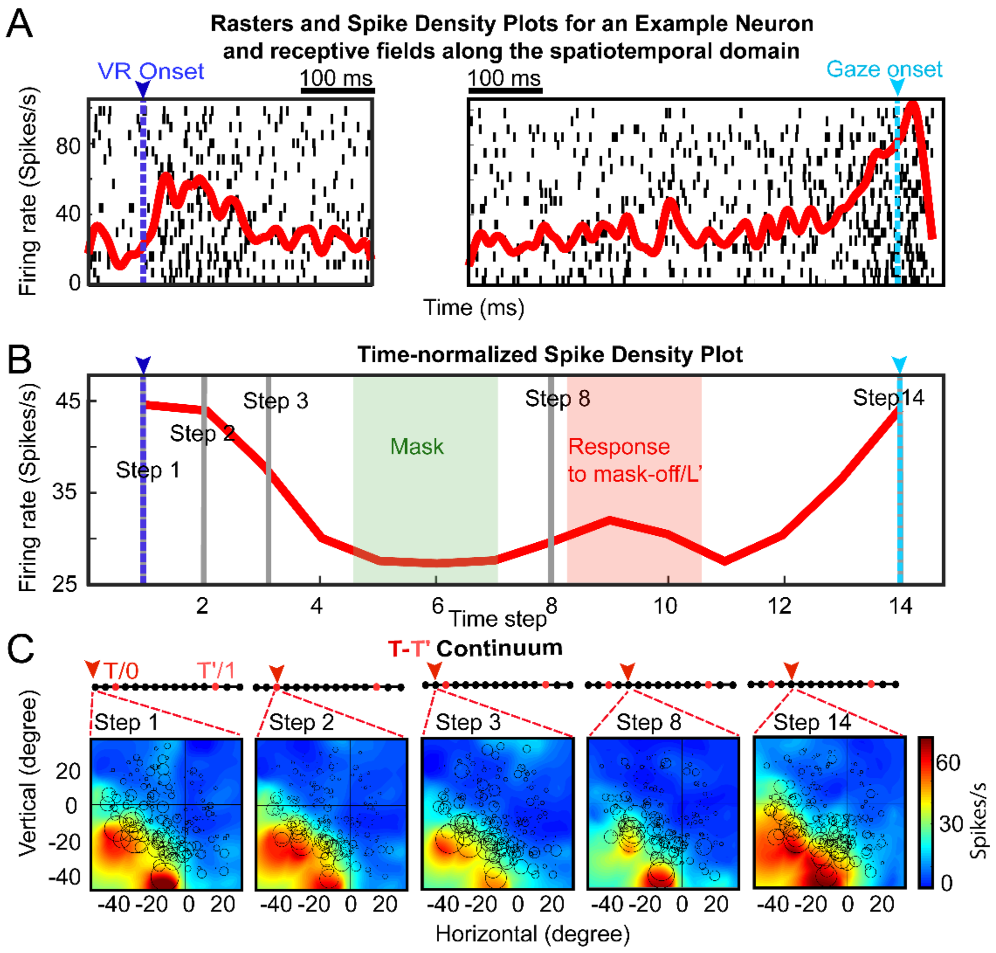


**Supplementary Fig. S9: An example of spatiotemporal analysis at the single cell level. (A)** Raster plot and spike density plot of a representative neuron with visual, delay and motor activity. To the left, the activity is aligned to the visual response onset (VR onset at 80 ms, blue line) and to the right, the activity is aligned to the gaze onset (cyan line). **(B)** Time-normalized activity (divided into 14 half-overlapping bins) of the raster in **'A'** aligned to VR onset (blue line) and the gaze onset (cyan line). The green shaded area corresponds to the mask duration and the red shaded area indicates the transient response to mask-off/landmark shift. **(C)** The RF maps of the neuron at different time-normalized steps (gray lines, corresponding to steps 1,2,3, 8 and 14) shown in **'B'**. The red arrowhead denotes the best fit model along the T-T' continuum (see top panel on each RF).The first three RFs include the VR burst and their RFs are fixed around T along the T-T' continuum, whereas at step 8 (after landmark shift) the RF shifts two steps toward T', then goes back to T (not shown) before finally shifting two steps toward T', just before the gaze onset. The colorbar indicates low-to-high activity from blue-to-red.
